# Cellular Substrate of Eligibility Traces

**DOI:** 10.1101/2023.06.29.547097

**Authors:** Léa Caya-Bissonnette, Richard Naud, Jean-Claude Béïque

## Abstract

The ability of synapses to undergo associative, activity-dependent weight changes constitutes a linchpin of current cellular models of learning and memory. It is, however, unclear whether canonical forms of Hebbian plasticity, which inherently detect correlations of cellular events occurring over short time scales, can solve the temporal credit assignment problem proper to learning driven by delayed behavioral outcomes. Recent evidence supports the existence of synaptic eligibility traces, a time decaying process that renders synapses momentarily eligible for a weight update by a delayed instructive signal. While eligibility traces offer a means of retrospective credit assignment, their material nature is unknown. Here, we combined whole-cell recordings with two-photon uncaging, calcium imaging and biophysical modeling to address this question. We observed and parameterized a form of behavioral timescale synaptic plasticity (BTSP) in layer 5 pyramidal neurons of mice prefrontal areas wherein the pairing of temporally separated pre- and postsynaptic events (0.5 s – 1 s), irrespective of order, induced synaptic potentiation. By imaging calcium in apical oblique dendrites, we reveal a short-term and associative plasticity of calcium dynamics (STAPCD) whose time-dependence mirrored the induction rules of BTSP. We identified a core set of molecular players that were essential for both STAPCD and BTSP and that, together with computational simulations, support a model wherein the dynamics of intracellular handling of calcium by the endoplasmic reticulum (ER) provides a latent memory trace of neural activity that instantiates synaptic weight updates upon a delayed instructive signal. By satisfying the requirements expected of eligibility traces, this mechanism accounts for how individual neurons can conjunctively bind cellular events that are separated by behaviorally relevant temporal delays, and thus offers a cellular model of reinforced learning.

## MAIN

Long-term synaptic plasticity is widely believed to contribute to the ability of the brain to learn ^1–4^. While the elucidation of the molecular and cellular underpinnings of synaptic plasticity mechanisms has received sustained attention over the last few decades, it remains unclear whether the canonical plasticity rules studied in experimentally tractable systems approximate well those guiding behaviorally relevant associative learning. For instance, while spike-timing-dependent plasticity (STDP) encapsulates key Hebbian postulates, it inherently relies on the correlation detection of cellular events occurring over time scales (tens of milliseconds ^5–9^) that are fundamentally distinct to those required for the processing of behaviorally relevant credit signal (at least hundreds of milliseconds ^1, 10^). As such, when instructive signals are delayed, a temporal credit assignment problem arises for which no unified solutions have yet emerged.

Eligibility traces are primarily a theoretical construct used in machine learning to implement credit assignment ^11^. In turn, they offer a useful heuristic to appraise synaptic models of associative encoding where they can be conceptualized as a silent process that keeps a time-decaying record of a synapse’s activity (*i.e.,* a short-lived local flag), rendering it temporarily ‘eligible’ for weight update upon a delayed instructive signal. Protracted eligibility traces would thereby allow individual neurons to solve the temporal credit assignment problem by binding cues that are separated by behaviorally relevant temporal delays. An emerging literature is beginning to identify forms of synaptic plasticity that exhibit temporal features that are consistent with the existence of a neural correlate of eligibility traces ^10, 12–16^. In particular, important recent work in the hippocampus has outlined the presence of a novel form of synaptic plasticity, termed Behavioral Timescales Synaptic Plasticity (BTSP), that emerges following the pairing of synaptic and cellular events that are separated by up to ∼ 2 seconds ^17–19^. While this form of plasticity was shown to be involved in the rapid emergence of place representation in CA1 neurons, the timescale of its induction rules alone provides direct evidence supporting the existence of eligibility traces in the brain. Yet, their material nature is unknown, and so are the details concerning the dynamical rules regulating their deployment.

Here, we show that the pairing of pre- and postsynaptic events with behaviorally relevant temporal delays induced synaptic potentiation in L5 pyramidal neurons of mice prefrontal areas (medial prefrontal cortex; mPFC) reminiscent of BTSP observed in the hippocampus. The protracted timescales of the plasticity induction preclude the involvement of canonical cellular mechanisms of plasticity. A series of two-photon (2P) glutamate uncaging and calcium (Ca^2+^) imaging experiments highlighted a spatially constrained, dendritic compartment specific, form of short-term associative plasticity of Ca^2+^ dynamics (termed here STAPCD) that emerged following the temporally discontiguous pairing of pre- and postsynaptic activity. Temporal parameterization, pharmacological experiments and computational modeling highlighted a mechanism wherein the endoplasmic reticulum (ER), through the concerted interplay between glutamate receptors of the N-methyl-D-aspartate receptor (NMDAR) and metabotropic glutamate receptor (mGluR) subtypes and ER Ca^2+^ channels of the ryanodine and inositol trisphosphate (IP_3_) subtypes, implement a delayed amplification of intracellular Ca^2+^. Blocking any of the core molecular players involved in STAPCD blocked BTSP, thereby pointing towards a causal relationship. These results thus describe a mechanism that holds a latent memory trace of cellular activity to participate in instantiating synaptic weight updates by a delayed instructive cue, and thus satisfies the features expected of eligibility traces.

## RESULTS

### BTSP in L5 pyramidal neuron of the mPFC

The experimental tractability of BTSP provides an opportunity not only to parametrize important dynamical features of eligibility traces, but also to determine their material nature. As an important corollary, we also sought to determine whether BTSP was solely operating in the hippocampus by examining whether qualitatively analogous protocols could induce plasticity in cortex. To achieve these goals, we prepared acute brain slices of the mPFC and carried out whole-cell recordings of layer 5 (L5) pyramidal neurons of male and female mice (Fig. 1A-B) using a potassium (K^+^)-Gluconate intracellular solution supplemented with picrotoxin (PTX), a γ-aminobutyric acid (GABA)_A_ receptor blocker, to isolate excitatory postsynaptic currents (EPSCs; see Fig. S1). We first examine the effects of a standard STDP protocol validated at other synapses ^9, 20, 21^ (*i.e.,* 120 pairings of a single synaptic input followed by 2 APs induced by a 50 ms current injection, with a 20 ms delay) and found, in keeping with previous reports ^12, 22^ that it did not induce significant potentiation of proximal synapses onto L5 PFC pyramidal neurons, even in young mice (P15 - P16; Fig. 1C; 96.71 ± 5.36 %) whose synapses are typically considered more plastic ^23^. Since bursting uncovers BTSP in hippocampus and regulates plasticity in cortex ^9, 17, 20, 21^, we next induced a presynaptic train of synaptic inputs (10 stimulations at 20 Hz; pre_t_) immediately prior to a burst of backpropagating action potentials (bAPs) induced by direct 300 ms current injection (post_b_), and found that it triggered robust potentiation in response to only 5 pairings (Fig. 1C; 123.96 ± 5.65 %). This burst-STDP (pre_t_-post_b_) was not restricted to younger animals as it was observed in mice P21 to P36 (Fig. S2A). Thus, a burst-STDP protocol not only induced potentiation at synapses that are otherwise resistant to potentiation by canonical STDP protocols, but did so following a far lower number of repetitions (5 *vs* 120).

**Figure 1.**
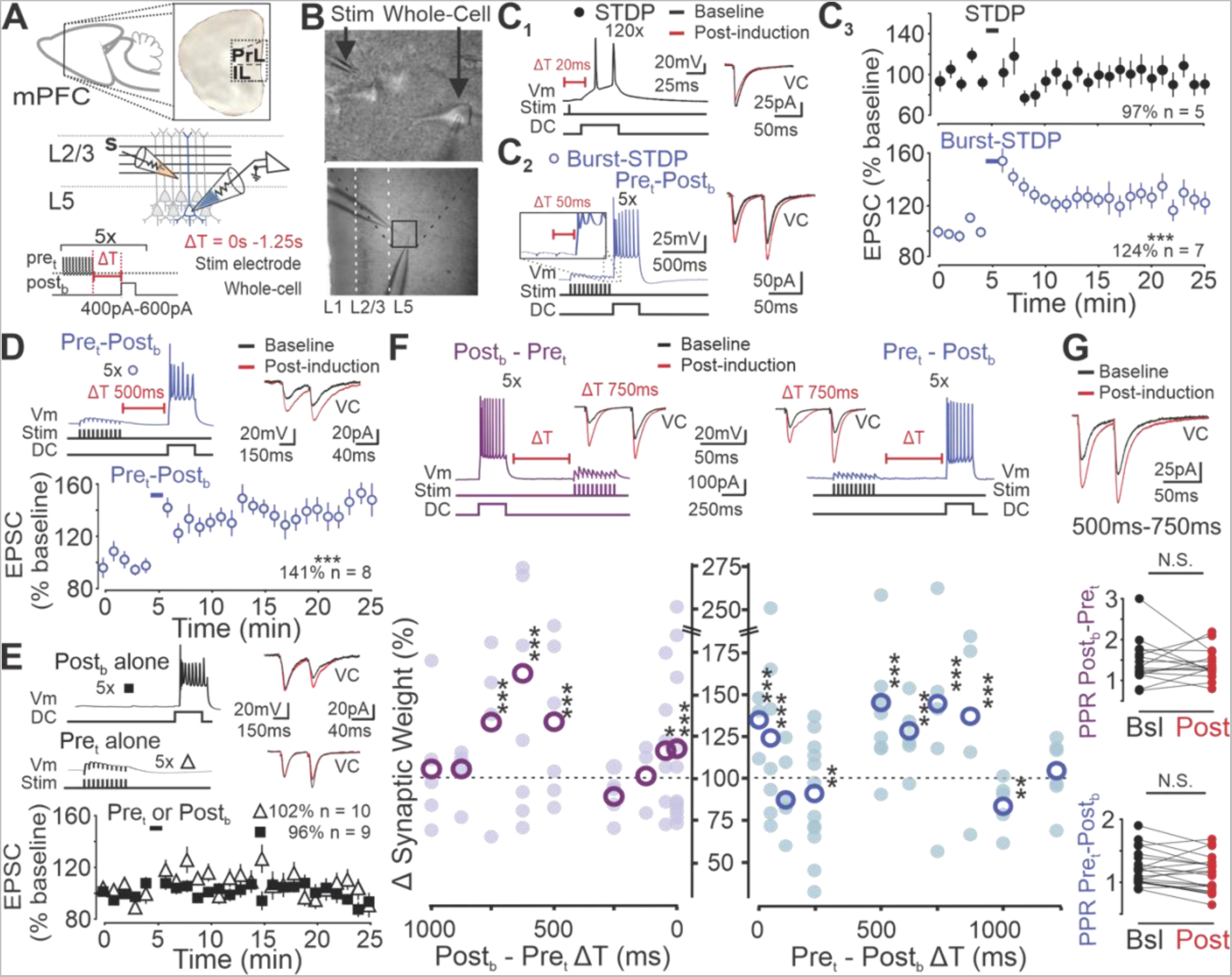
Synapses onto mPFC L5 pyramidal neurons undergo BTSP. (A) Schematic of experimental approach: a stimulating electrode (s) is placed proximal to L5 neuron apical dendrites and a recording electrode in whole-cell patch clamp records from a L5 pyramidal neuron (PrL; prelimbic, IL; infralimbic). (B) Example image of L5 pyramidal neurons of the mPFC using 40 X submersive lens (top), and 5 X lens (bottom). (C) C_1_ Representative trace of STDP pairings in current clamp (left) and baseline and post-pairing EPSCs in voltage clamp (VC; right). C_2_ Same as in C_1_ except for burst-STDP pairings. C_3_ Normalized EPSC changes over 5 minutes of baseline and 20 minutes of post-induction recordings for STDP induction (top: n = 5 neurons, p = 0.71) and burst-STDP induction (bottom: n = 7 neurons, p = 0.0007) protocols induced at the time indicated by color coded bars. (D) Representative trace of pre_t_- post_b_ pairings at 500 ms delay and EPSCs (top). Resulting normalized EPSC change over time (bottom, n = 8 neurons, p = 1.07 x 10^-7^). (E) Representative trace of post_b_ (top, squares) or pre_t_ (bottom, triangles) induced alone (top left) and EPSCs (top right). Resulting normalized EPSC change over time (bottom, post_b_: n = 9 neurons, p = 0.43, pre_t_: n = 10 neurons, p = 0.88). (F) Representative trace of the pre_t_-post_b_ pairings and post_b_-pre_t_ pairings and EPSCs (top). Resulting mean synaptic weight change induced at different timing intervals from 0 ms to 1.25 s (pre_t_-post_b_ 0 ms: n = 3 neurons, p = 1.03 x 10^-7^, 50 ms: as shown in C_3_, 125 ms: n = 5 neurons, p = 0.23, 250 ms: n = 12 neurons, p = 0.007, 500 ms: as shown in F, 625 ms: n = 7 neurons, p = 5.04 x 10^-7^, 750 ms: n = 5 neurons, p = 0.0002, 875 ms: n = 4 neurons, p = 7.12 x 10^-5^, 1000 ms: n = 5 neurons, p = 0.001, 1250 ms: n = 6 neurons, p = 0.23; post_b_-pre_t_ 0 ms: n = 12 neurons, p = 7.46 x 10^-5^, 50 ms: n = 6 neurons, p = 0.01, 125 ms: n = 3 neurons, p = 0.95, 250 ms: n = 5 neurons, p = 0.054, 500 ms: n = 6 neurons, p = 7.20 x 10^-6^, 625 ms: n = 7 neurons, p = 9.18 x 10^-7^, 750 ms: n = 4 neurons, p = 0.0008, 875 ms: n = 4 neurons, p = 0.42, 1000 ms: n = 5 neurons, p = 0.24). (G) Representative trace of baseline and post-induction EPSCs (top). PPR for protocols shown in G at time intervals between 500 ms and 750 ms (middle: post_b_-pre_t_, n = 16 neurons, p = 0.73, bottom: pre_t_-post_b_, n = 20 neurons, p = 0.054). bsl; baseline. ***p < 0.001, ** p < 0.01, *p < 0.05; Data are shown as mean ± sem.

We then asked whether bursting in L5 pyramidal neurons would bend the canonical temporal rules of STDP induction into the realm of BTSP (*i.e.,* delays in the hundreds of milliseconds). We thus extended the temporal delay between presynaptic and postsynaptic stimulations to 500 ms, a timescale far outlasting that of STDP induction ^6–8, 24, 25^, and found, strikingly, that this protocol induced potentiation (Fig. 1D; 140.87 ± 5.43 %). Importantly, neither presynaptic trains alone, nor postsynaptic bursts alone, yielded potentiation (Fig. 1E), confirming that this form of plasticity at extended timescale is associative in nature.

To investigate with greater granularity the induction rules of BTSP in L5 neurons, we parameterized the timing delay between pre- and postsynaptic events (0 s – 1.25 s). While the pre_t_-post_b_ protocol induced a significant potentiation at short- (0 ms - 50 ms) and extended- (500 ms - 900 ms) time intervals, it induced a small but highly reliable depression of synaptic strength at intermediate timings (125 ms - 250 ms; Fig. 1F). Since the polarity of the synaptic changes induced by canonical STDP is determined by the relative order of pre- and postsynaptic spiking ^6, 7^, we naturally next investigated the effects of reversing the order of pre- and postsynaptic firing. The temporal profiles of both induction protocols (*i.e.,* pre_t_-post_b_ and post_b_-pre_t_) were remarkably symmetrical, with both showing strong potentiation at short and extended timescales, and depression at intermediate delays (Fig. 1F). We next explored induction singularities of BTSP at timescales that induced potentiation (*i.e.,* 500 ms - 900 ms). We found that the magnitude of BTSP showed a small positive correlation (pearson correlation coefficient = 0.34, p = 0.04) with the magnitude of short-term facilitation of the stimulated synapses (Fig. S2B). It was otherwise not dependent on a range of experimental variables, including age and sex of the animals, as well as Ih currents and spike frequency adaptation (see Fig. S2), implying a generalization and robustness of these plasticity rules across L5 pyramidal neuron subtypes ^26, 27^. The paired-pulse ratio (PPR) of baseline *vs* post-pairing EPSCs, an index of presynaptic release probability ^28^, did not change significantly, suggesting a postsynaptic locus of expression (Fig. 1G).

**Figure 2.**
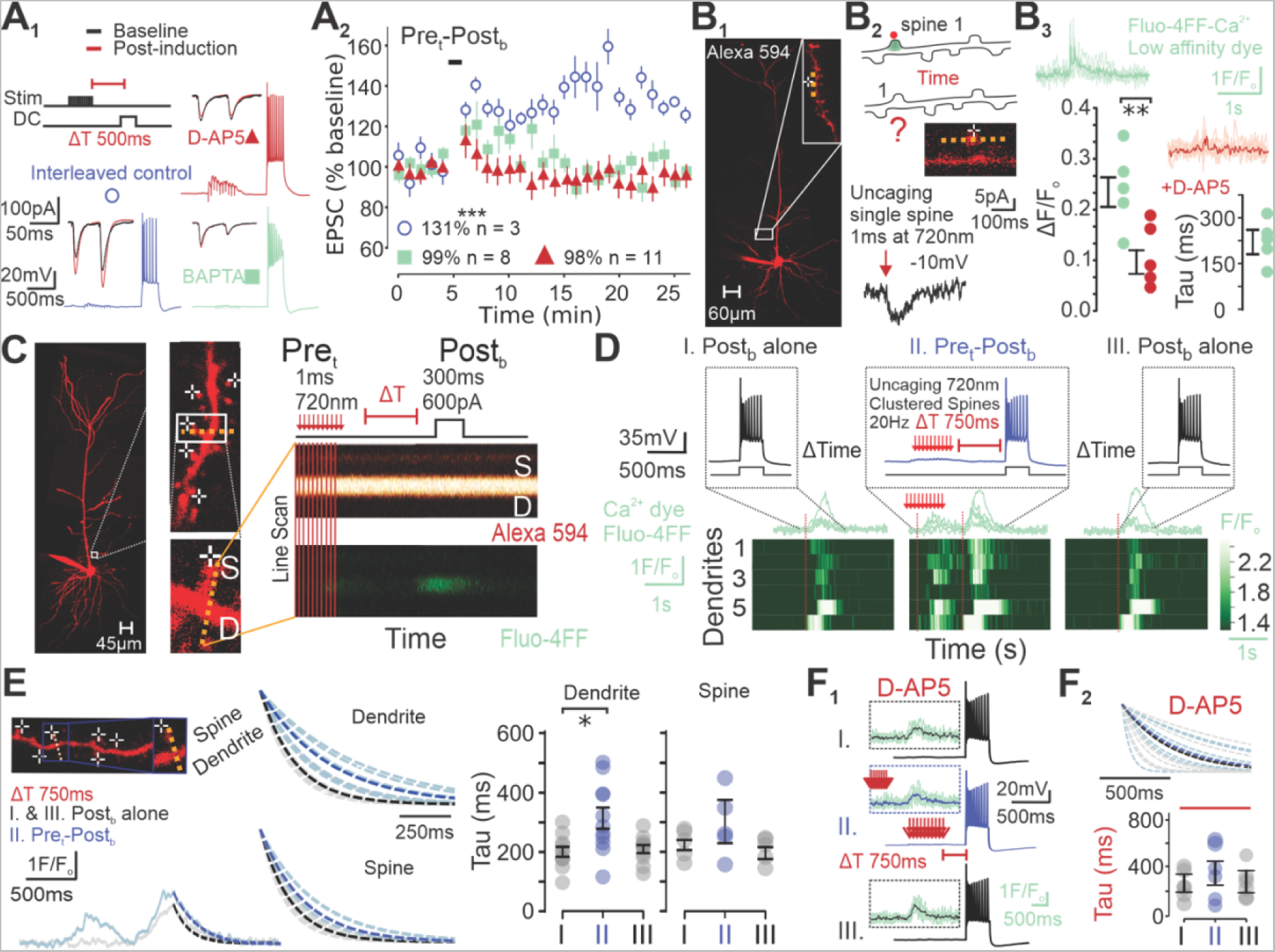
Synaptically-driven short-term associative plasticity of Ca^2+^ dynamics. (A) A_1_ Representative trace of pre_t_-post_b_ BTSP induction for interleaved control (blue), or with bath- application of D-AP5 (red) or intracellular BAPTA (green), and respective baseline and post- induction EPSCs. A_2_ Normalized EPSC change over time (blue: n = 3 neurons, p = 1.07 x 10^-9^, red: n = 11 neurons, p = 0.36, green: n = 8 neurons, p = 0.66). (B) B_1_ Example image of L5 pyramidal neuron and proximal apical dendrite for which glutamate uncaging and line scans were performed at the spine. B_2_ Cartoon of experiment (top). Example image of proximal apical spine (middle). Representative trace of 1 EPSC induced by glutamate uncaging at -10 mV (bottom). B_3_ Ca^2+^ traces obtained with Fluo-4FF dye for protocol shown in B_2_ in control condition (green) and following bath-application of D-AP5 (red) (n = 5 neurons; top) and resulting ΔF/F (p = 0.0063) and Tau values (bottom). (C) Example of 2P line scans recording of a L5 proximal apical dendrite during uPre_t_-post_b_ pairing using Fluo-4FF dye. (D) Example traces of three post_b_ given at 30 s intervals with a uPre_t_ given 750 ms prior to the second post_b_, and resulting Ca^2+^ traces measured by line scans using Fluo-4FF (n = 6 dendrites). (E) Resulting decay traces and tau values of post_b_ from dendritic line scans as shown in D (n = 11 dendrites, p = 0.024) and at the spines (n = 5 spines, p = 0.19). (F) F_1_ Representative voltage traces and resulting Ca^2+^ traces of protocol shown in D, except in the presence of D-AP5. F_2_ Resulting decay traces and tau values (n = 6 dendrites, p = 0.44). *p < 0.05, **p < 0.01; Data are shown as mean ± sem.

Collectively, these results show that L5 neurons can bind pre- and postsynaptic events, presented in indiscriminate order, that are separated by time intervals far longer than those of canonical STDP and of typical membrane time constants ^6, 25^. The timescale and symmetry of this form of BTSP makes it non-Hebbian in nature and represent a direct manifestation of protracted synaptic eligibility traces in L5 pyramidal neurons.

### Short-Term Plasticity of Ca^2+^ Dynamics

While eligibility traces have provided a useful framework to formalize several computational implementations of plasticity ^10, 12, 29^, their material nature has not been established. In an attempt to address this knowledge gap, we first tested the general role of postsynaptic Ca^2+^ in BTSP and found that pre_t_-post_b_ BTSP was abolished by the fast Ca^2+^ buffer BAPTA (20 mM) as well as by bath administration of the NMDAR antagonist D-AP5 (100 µM; Fig. 2A). Since mPFC synapses contain a high proportion of high affinity, slow, GluN2B- containing NMDARs ^30, 31^, we wondered whether singularly protracted NMDAR-dependent Ca^2+^ influx may underlie the extended coincident detection between pre- and postsynaptic bursting of BTSP. We thus carried out simultaneous 2P uncaging of MNI-glutamate and Ca^2+^ imaging of the low affinity dye Fluo-4FF. These experiments revealed that the decay of Ca^2+^ transients (218 ms ± 41 ms) on single spines of proximal apical dendrites of L5 pyramidal neurons were: 1) broadly similar to those observed at canonical mature CA1 synapses ^31–34^, and; 2) significantly faster than the time interval of effective BTSP with pre_t_-post_b_ protocols (Fig. 2B). In an attempt to more closely mimic our pre_t_-post_b_ BTSP experiments, we also assessed the decay of Ca^2+^ transients following the repetitive synaptic stimulation (10 uncaging stimulations at 20 Hz) of spatially clustered spines (5 - 6 spines) in current clamp mode (uPre_t_; Fig. S3A) and found that Ca^2+^ transients were still substantially shorter than the typical delay of paired inputs during BTSP (Fig. S3B). Given that the decay of fluorescence, even when using a low affinity dye, likely overestimates the lifetime of phasic cytosolic Ca^2+^ *in situ* ^33, 35^, these results suggest that the cytosolic Ca^2+^ triggered by synaptic activation onto L5 pyramidal neurons has largely subsided by the time the post_b_ invades the dendritic arbor during effective BTSP induction, especially for the longer intervals (500 ms to 1000 ms). A protracted, synaptically-induced, cytosolic Ca^2+^ trace is therefore an unlikely candidate for enacting an eligibility trace in L5 mPFC neurons.

For canonical forms of Hebbian plasticity, much of the scrutiny has historically focused on the dynamics of NMDAR-dependent Ca^2+^ entry occurring during the ephemeral coincident detection phase of plasticity induction ^32, 33, 36^. Here, the Ca^2+^ entering during the delayed instructive event (either synaptic or bAPs) may, in principle, act as the plasticity signaling molecule permissive to synaptic potentiation during BTSP. We thus re-focused our attention and examined features of the Ca^2+^ transients triggered by the delayed instructive postsynaptic burst occurring during pre_t_-post_b_ BTSP induction. Using 2P Ca^2+^ imaging, we first established the Ca^2+^ profile at proximal apical dendrites and associated spines induced by a burst of bAPs alone (post_b_ alone; denominated as ‘I.’ in Fig 2-5). We restricted our experiments to primary oblique dendrites and did not observe branch point failures of bAP propagation (at least as inferred from Ca^2+^ imaging). Following a 30 second rest period, we re-imaged the same dendritic segment in response to post_b_, but when it was *preceded* by synaptic activation (by 750 ms), achieved here by glutamate uncaging onto a cluster of closely positioned spines (uPre_t_) (mimicking pre_t_-post_b_ BTSP induction; denoted ‘II.’ in Fig. 2-5). Remarkably, we found that the decay time constant of dendritic Ca^2+^ triggered by a burst of bAPs (post_b_) was nearly doubled when the burst of bAPs were preceded by clustered synaptic activation (compared to post_b_ alone; II. *vs* I.; Fig. 2C-E). This robust augmentation of Ca^2+^ transients was manifest in the dendrites, but not in the stimulated spines, suggesting a dendritic origin (Fig. 2E). When probed following another 30 second rest period, the decay of the Ca^2+^ transients induced by post_b_ alone (III.) typically returned to that of the initial isolated post_b_ (III. *vs* I.; Fig. 2D-E, Fig. S4), suggesting that the cellular phenomenon supporting this synaptically-induced prolongation of Ca^2+^ signals induced by bAPs is short-lived. We have formalized this general observation as an exclusion criterion to restrict our analysis to healthy recordings (*i.e.,* where the baseline value (I.) and post-paring value (III.) were within 35 % of each other; see Methods). Not surprisingly, the administration of D-AP5 blocked the uncaging-induced synaptic Ca^2+^ transients (Fig. 2B) and did not influence the profile of the Ca^2+^ transients induced by post_b_ alone. However, it blocked the lengthening of bAPs burst-triggered Ca^2+^ transients induced by preceding synaptic activation (Fig. 2F; Fig. S4). Thus, these results imply that the Ca^2+^ transients triggered by bursts of bAPs are increased when preceded (750 ms before) by a bout of synaptic activation, which effect is dependent on NMDARs. The plasticity of Ca^2+^ transients represents a plausible and intriguing mechanism for enacting an eligibility trace for binding pre- and postsynaptic inputs at behavioral timescales. For simplicity and ease, we will heretofore refer to this phenomenon as Short-Term Associative Plasticity of Ca^2+^ Dynamics (STAPCD).

**Figure 4.**
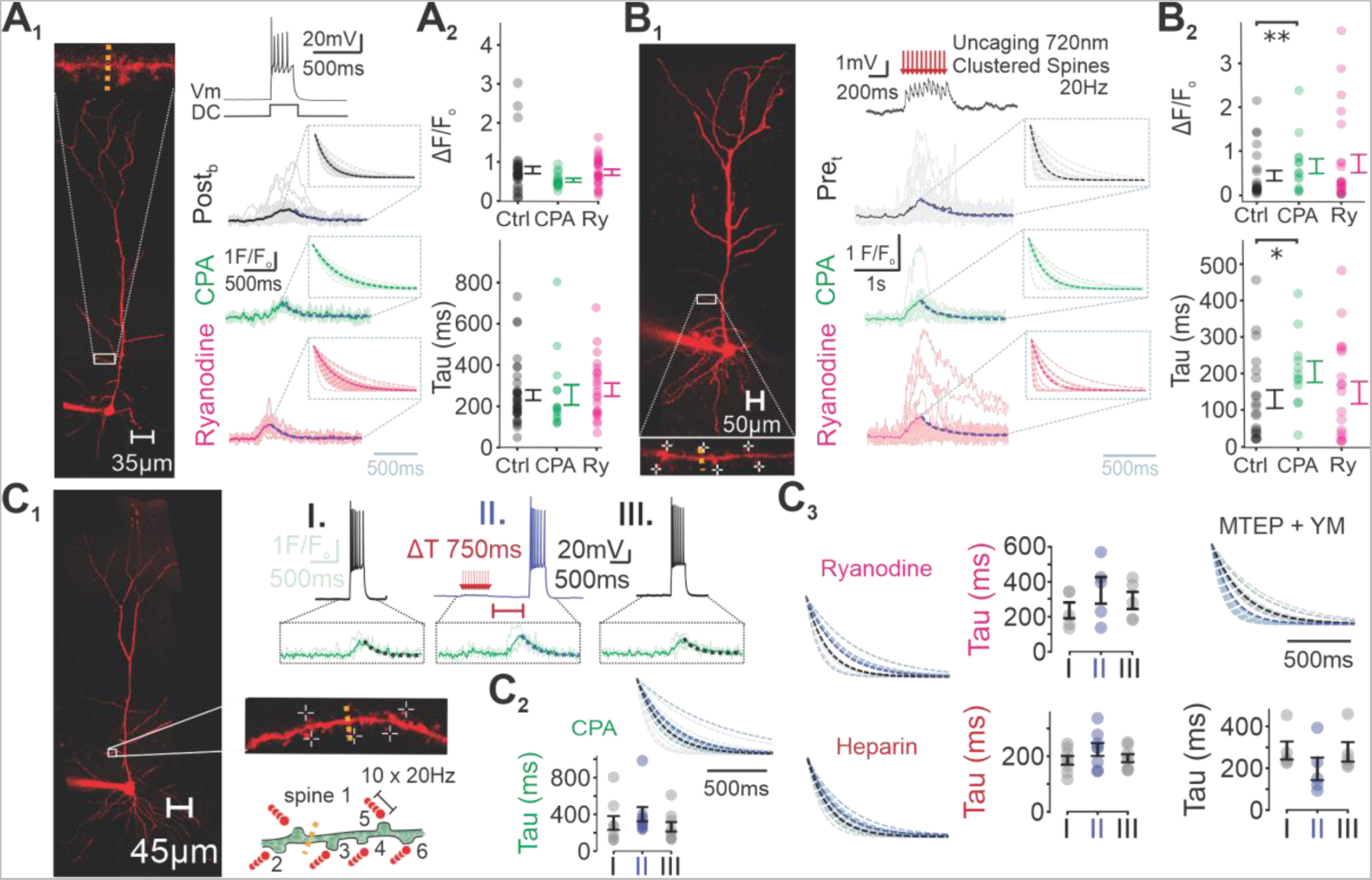
Ca^2+^ stores sustain STAPCD. (A) A_1_ Representative voltage trace, and resulting Ca^2+^ and decay traces of post_b_ performed on L5 proximal apical dendrites in control condition (black, n = 38 dendrites), with bath-application of CPA (green), or intracellular ryanodine (pink). A_2_ Resulting peak Ca^2+^ fluorescence (CPA: n = 14 dendrites, p = 0.40, ryanodine: n = 24 dendrites, p = 0.18) and decay time constant (CPA: n = 14 dendrites, p = 0.42, ryanodine: n = 22 dendrites, p = 0.19). (B) Same as A except for uPre_t_ ΔF/F (control: n = 38 dendrites, CPA: n = 14 dendrites, p = 0.0077, ryanodine: n = 24 dendrites, p = 0.086), and decay time constant (control: n = 22 dendrites, CPA: n = 12 dendrites, p = 0.016, ryanodine: n = 20 dendrites, p = 0.49). (C) C_1_ Representative voltage trace and resulting Ca^2+^ traces of three post_b_ given at 30 s intervals with the second post_b_ preceded by uPre_t_ 750 ms prior in the presence of bath-applied drugs. C_2_ Resulting decay traces and time constants in the presence of extracellular CPA (n = 8 dendrites, p = 0.15), C_3_ Same as C_2_ except for intracellular ryanodine (pink: n = 7 dendrites, p = 0.13), intracellular heparin (red: n = 8 dendrites, p = 0.15), or extracellular mGluR antagonists, MTEP and YM-298198 (black: n = 5 dendrites, p = 0.063; bottom). *p < 0.05, **p < 0.01. Data are shown as mean ± sem.

**Figure 5.**
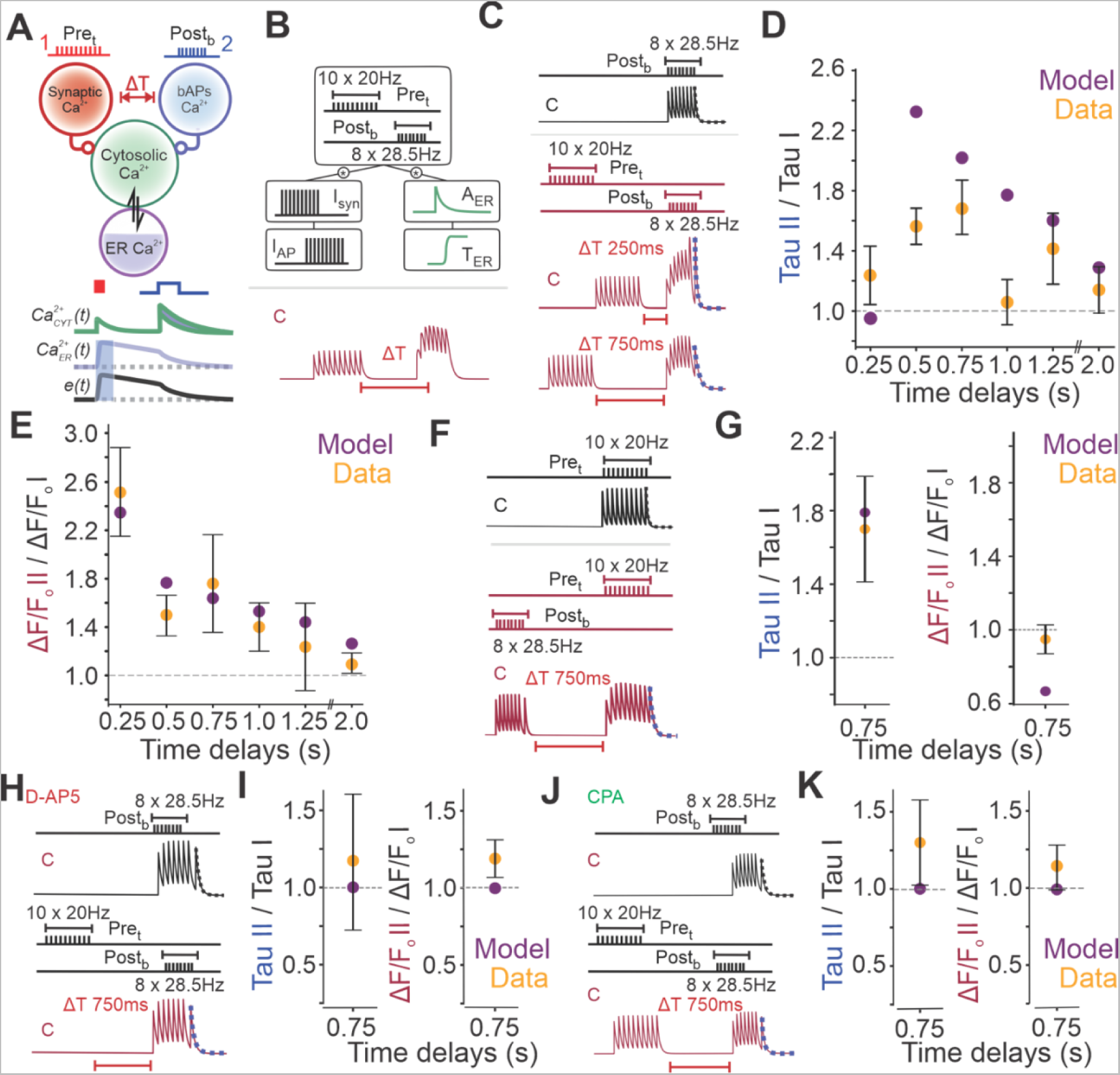
Plasticity of Ca^2+^ dynamics are captured by combining synaptic and bAPs Ca^2+^ with ER modulation. (A) Schematic depicting the relationship between synaptic, bAPs, cytosolic and ER Ca^2+^, and with eligibility traces. (B) Schematic depicting the nonlinear Ca^2+^ dynamics fitted by minimizing least-squares approach. (C) Modeled cytosolic Ca^2+^ traces of post_b_ (black) and pre_t_-post_b_ pairings at 250 ms and 750 ms (red). (D) Relative normalized decay time constants of post_b_ paired with pre_t_ compared to post_b_ induced alone at varying time delays. (E) Same as D except for peak amplitude of Ca^2+^ transient. (F) Modeled Ca^2+^ traces of pre_t_ (black) and post_b_-pre_t_ pairings at 750ms (red). (G) Relative normalized decay time constants (left) and peak amplitude (right) of traces shown in F. (H) Modeled Ca^2+^ traces of post_b_ (black) and pre_t_- post_b_ pairings at 750ms (red) with D-AP5 (I_syn_ is set to 0). (I) Relative normalized decay time constants (left) and peak amplitude (right) of traces shown in H. (J) Modeled Ca^2+^ traces of post_b_ (black) and pre_t_-post_b_ pairings at 750ms (red) with CPA (A_ER_ and T_ER_ are set to 0). (K) Relative normalized decay time constants (left) and peak amplitude (right) of traces shown in J. Data are shown as mean ± sem.

### Induction Rules of STAPCD abide by those of BTSP

As a minimal necessary condition for a causal relationship inference between STAPCD and BTSP, their temporal induction rules should be broadly similar. We therefore investigated with increased granularity the temporal contingencies of the delays between the synaptic activation and the burst of bAPs (0.25 s - 4.5 s) and found that STAPCD was readily expressed following extended intervals (0.5 s - 1 s), but not at 250 ms, a temporal profile closely matching BTSP induction in these neurons (Fig. 3A). STAPCD was also expressed as an amplification in the amplitude (ΔF/F) *per se* of the Ca^2+^ signal for a temporal window of up to ∼ 1 s (Fig. S4; see Ca^2+^ model below). We next examined the bidirectionality of STAPCD by determining whether, like BTSP, it was induced when a burst of bAPs (post_b_) preceded synaptic activation (post_b_-uPre_t_). We thus established the baseline profile of synaptic Ca^2+^ transients (again induced by 2P uncaging of MNI-glutamate onto visually identified clustered spines; uPre_t_), and repeated the procedure 30 seconds later but with the uncaging stimulation (uPre_t_) preceded by burst of bAPs (post_b_; 750 ms before). The decay of Ca^2+^ transients induced by the uncaging trains that were preceded by bursts of bAPs were significantly longer than those triggered by synaptic trains alone (*i.e.,* baseline conditions measured 30 seconds before and after the post_b_- uPre_t_ pairing; Fig. 3B; Fig. S5A-B). Altogether, the bidirectionality and the temporal requirements of STAPCD closely matches those of BTSP.

**Figure 3.**
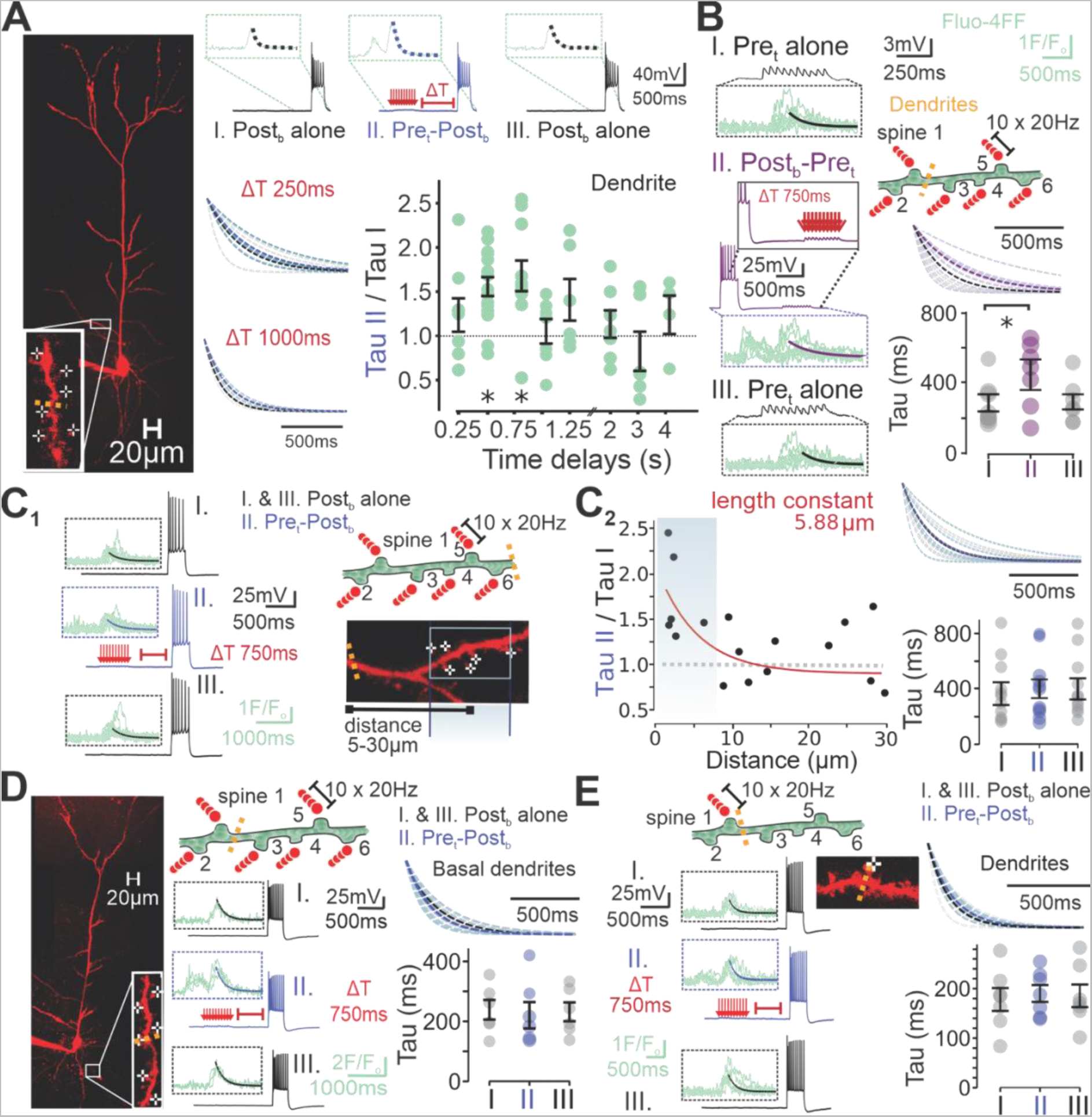
Properties of STAPCD during effective BTSP. (A) Representative voltage and Ca^2+^ trace of three post_b_ given at 30 s intervals with a uPre_t_ given at varying time intervals prior to the second post_b_ (top), and resulting decay traces for time intervals of 250 ms and 1 s (middle). Normalized relative decay time constants of post_b_ preceded by uPre_t_ (II.) compared to the first post_b_ given alone (I.; 250 ms: n = 8 dendrites, p = 0.31, 500 ms: n = 18 dendrites, p = 0.030, 750 ms: as shown in Fig. 2E, 1000 ms: n = 7 dendrites, p = 0.81, 1250 ms: n = 6 dendrites, p = 0.31, 2000 ms: n = 7 dendrites, p = 0.81, 3000 ms: n = 6 dendrites, p = 0.69, 4500 ms: n = 4 dendrites, p = 1.0) (bottom right). (B) Representative voltage and Ca^2+^ traces of three uPre_t_ given at 30 s intervals with the second uPre_t_ preceded by post_b_ 750 ms prior induced on L5 proximal apical dendrites (left). Resulting decay traces and time constants (right: n = 7 dendrites, p = 0.031). (C) C_1_ Experimental setup (right), and representative voltage trace and resulting Ca^2+^ traces of three post_b_ given at 30 s intervals with the second post_b_ preceded by uPre_t_ 750 ms prior (left). C_2_ Distance dependence of normalized relative decay time constants. Length constant was measured by mono-exponential fit (left). Resulting decay traces and time constants for line scans performed 5-30 µm away from the centroid of clustered spine activation (right, n = 11 dendrites, p = 0.58). (D) Example image of L5 pyramidal neuron and a proximal basal dendrite (left). Representative cartoon, voltage trace and resulting Ca^2+^ traces measured on basal dendrites during three post_b_ given at 30 s intervals with the second post_b_ preceded by uPre_t_ 750 ms prior (middle), and resulting decay traces and time constants (right: n = 6 dendrites, p = 0.56). (E) Experimental setup (top left). Representative voltage trace and resulting Ca^2+^ traces of three post_b_ given at 30 s intervals with the second post_b_ preceded by uPre_t_ given on a single spine 750 ms prior (bottom left). Resulting decay traces and time constants (right: n = 7 dendrites, p = 0.81). * p < 0.05, Data are shown as mean ± sem.

We next parameterized some of the salient features of STAPCD. First, we determined its spatial profile and found that the synaptically-induced amplification of bAPs burst-induced Ca^2+^ transients was largely restricted to the dendritic segment of the activated spine cluster (average 7.83 µm; n = 17 dendrites), sharply decreasing with increasing distance along the dendrite (length constant of ∼6 µm from cluster centroid; Fig. 3C; Fig. S6A). Second, this associative form of Ca^2+^ plasticity showed cellular compartment specificity: it was readily observed at proximal apical oblique dendrites but not at basal dendrites (Fig. 3D, Fig. S6B). Third, we observed that STAPCD depended on cooperativity between spatially clustered spines since train stimulations of a single spine did not significantly alter the Ca^2+^ transients induced by bursts of bAPs (Fig. 3E; Fig. S6C). This result suggests that the manifestation of STAPCD rely on a minimal amount of Ca^2+^ influx during the synaptic cue. Consistent with this idea, the magnitude of STAPCD was slightly correlated to the amplitude of the initial Ca^2+^ transients induced by clustered synaptic activation (uPre_t_; Fig. S7D-E). The extent of STAPCD appeared to be somewhat bounded since the Ca^2+^ amplification was of a lesser magnitude in cases where the bAPs displayed longer Ca^2+^ transients to begin with (i.e., ‘I”; Fig. S7B).

We finally report a range of experimental parameters that were not correlated with the manifestation of STAPCD (see Fig. S7). Collectively, these results outline a set of core features of a novel form of intracellular Ca^2+^ plasticity. We found that STAPCD: 1) is triggered by the conjunctive actions of synaptic and bAP events occurring over protracted time scales, 2) is spatially compartmentalized, and; 3) showed induction rules that closely parallel those of BTSP.

### Intracellular Ca^2+^ Stores are involved in the expression of STAPCD

Pondering on the temporal and associative features of the intracellular Ca^2+^ plasticity outlined above, we conjectured that intracellular Ca^2+^ stores may be involved in its manifestation. Indeed, the smooth ER is involved in intracellular Ca^2+^ homeostasis, is found as an anastomosing network in dendrites of pyramidal neurons ^37–39^ and, at times, in spines ^38, 40^. The ER regulates intracellular Ca^2+^ dynamics through seemingly opposing uptake and release mechanisms (*i.e.,* involving either Ca^2+^ uptake/extrusion from, or release into, the cytoplasm) ^41–44^. In neurons, ER-mediated mechanisms have been shown to regulate in some conditions the profile of both synaptically-induced and spiking-induced cytosolic Ca^2+^ entry ^32, 45–47^, and to be involved in several forms of synaptic plasticity ^32, 48^. In order to ultimately determine whether ER Ca^2+^ stores are involved in STAPCD, we first began by determining the contribution of ER stores to the profile of Ca^2+^ induced by either synaptic trains or bursts of bAPs alone in L5 mPFC neurons. We thus incubated slices in CPA, a Sarco/Endoplasmic Reticulum Ca^2+^-ATPase (SERCA) pump blocker, for at least 1 h prior to recordings. While CPA did not alter the Ca^2+^ dynamics induced by post_b_, it modestly, but significantly, increased the decay time constant of dendritic Ca^2+^ profile induced by trains of 2P synaptic activation (uPre_t_) (Fig. 4A-B). These results suggest that SERCA pumps uptake cytosolic Ca^2+^ that enters following trains of glutamate release and thus slightly constrains its cytosolic temporal profile in proximal apical dendrites of L5 pyramidal neurons in baseline conditions. Next, given that ER stores can in certain conditions amplify cytosolic Ca^2+^ by a process known as Ca^2+^-induced Ca^2+^-release (CICR), we examined the effects of blocking CICR by including ryanodine in the intracellular recording solution. In our conditions, the temporal profile of dendritic Ca^2+^ elicited by either burst of bAPs or synaptic inputs were not altered by ryanodine (Fig. 4A-B), thus suggesting that ER stores do not appreciably sustain CICR in response to isolated pre- or post- synaptic events.

An intuition that naturally emerges from the minimal contribution of CICR to synaptic and bursts of bAPs events in baseline conditions, along with the Ca^2+^ uptake by SERCA pumps, is that the ER stores may exhibit a conditional, state-dependent, CICR capacity that would manifest itself following temporally paired Ca^2+^ events. To begin addressing this possibility, we determined the effects of blocking the SERCA pumps on STAPCD and found that CPA prevented it (uPre_t_ 750 ms before post_b_; Fig. 4C; Fig. S8). In line with a role of CICR, intracellular ryanodine significantly reduced STAPCD, but ostensibly only partially (Fig. 4C; Fig. S8). Thus, the expression of STAPCD requires intact ER stores through a mechanism that, at least partially, involves CICR.

In order to delineate more precisely the involvement of ER in STAPCD, we wondered whether the partial block by ryanodine may reflect the parallel involvement of IP_3_-induced Ca^2+^-Release (IICR) ^41^, especially given the second-long timescale of action of IP_3_ ^42, 49^. We thus blocked IP_3_ receptors (IP_3_Rs) with intracellular heparin and found that it abolished STAPCD (Fig. 4C; Fig. S8). The involvement of IP_3_Rs was unsuspected in that it suggests the involvement of a Gq-coupled receptor signaling pathway ^50, 51^. As a first candidate, we blocked mGluRs of the mGluR1/5 subtype with bath-administration of MTEP and YM-298198, and found that it abolished STAPCD (Fig. 4C; Fig. S8). Collectively, these results thus far demonstrate that STAPCD requires activation of glutamate receptors of the NMDAR and mGluR subtypes, as well as CICR triggered by activation of ryanodine receptors (RyRs) and IP_3_Rs on intact ER stores.

### Activity-dependent state transition of ER Ca^2+^ stores accounts for STAPCD

While the results outlined above identified a set of core molecular players involved in STAPCD, they however fall short of providing a dynamical and mechanistic account of STAPCD. We thus turned to computational simulations to determine whether the salient features of STAPCD could be modeled by a parsimonious set of experimentally grounded parameters. In particular, we sought to capture the observations that suggest that while ER stores, in baseline conditions, primarily uptake (as opposed to release) Ca^2+^, they can conditionally amplify the Ca^2+^ transient induced by a delayed signal. We thus modeled cytosolic Ca^2+^ (*c*) as reflecting direct entry arising from synaptic (*I_syn_*) and bAPs (*I_AP_*) sources, as well as indirect contribution by intracellular Ca^2+^ stores through cytosolic uptake/extrusion (ER loading) and release (ER unloading) mechanisms (Fig. 5A-B; see Methods for details). The ER amplification of cytosolic Ca^2+^ (*i.e.,* ER unloading) by a delayed instructive cue is temporally constrained to a period of ER release eligibility (*i.e.,* ∼ 1 s; experimentally determined in Fig 3A) and consists in a linear modification in cytosolic Ca^2+^ amplitude (*A_ER_*) and a non-linear modification in Ca^2+^ decay time constant (*T_ER_*). *A_ER_* reflects Ca^2+^ release through IICR and CICR, while the non-linearity of *T_ER_* reflects a previously reported concentration-dependent non-linear rate of Ca^2+^ transport by SERCA pumps ^43, 44^.

Simulations deployed from our model captured the temporal rules of STAPCD induction (Fig. 5C-G; Table S1) that were determined experimentally. We further simulated the effects of blocking NMDARs with D-AP5 by setting *I_syn_* to 0, and the model reproduced our experimental data in amplitude and decay time constant (Fig. 5H-I). To simulate our experiments blocking SERCA pumps with CPA, we have set both the *A_ER_* and *T_ER_* parameters to 0, and the model once again reproduced the experimental data (Fig. 5J-K). Together, our experiments and model simulations suggest a mechanistic scenario underlying STAPCD: Ca^2+^ entry triggered by synaptic inputs or postsynaptic firing alters the state of the ER by rendering it momentarily (*i.e.,* ∼ 1 second) eligible to sustain RyR- and IP_3_R-mediated intracellular Ca^2+^release, that is triggered by a secondary, time delayed, Ca^2+^ entry associated with an instructive cue. As such, the ER effectively retains a memory of previous activity on a second-long timescale, yielding STAPCD.

### Causal relationship between STAPCD and BTSP

The results from our pharmacological investigation and simulations identified a set of core molecular players involved in STAPCD along with a plausible mechanistic and dynamical model for the role of the ER in STAPCD. In turn, this level of description provides an opportunity to explore the causal relationship between STAPCD and BTSP and, as a corollary, whether STAPCD enact an eligibility trace in cortex. As an obligatory requirement for a causal relationship, the pharmacological blockade of every molecular component known to be involved in STAPCD in isolation, should abolish BTSP. We tested this conjecture and found that individually blocking: 1) NMDARs (with D-AP5; Fig. 2A); 2) SERCA pumps (with CPA; Fig. 6A); 3) RyRs (with intracellular ryanodine; Fig. 6B); 4) IP_3_Rs (with intracellular heparin; Fig. 6B) or; 5) mGluRs (with MTEP and YM-298198; Fig. 6B) abolished BTSP.

**Figure 6.**
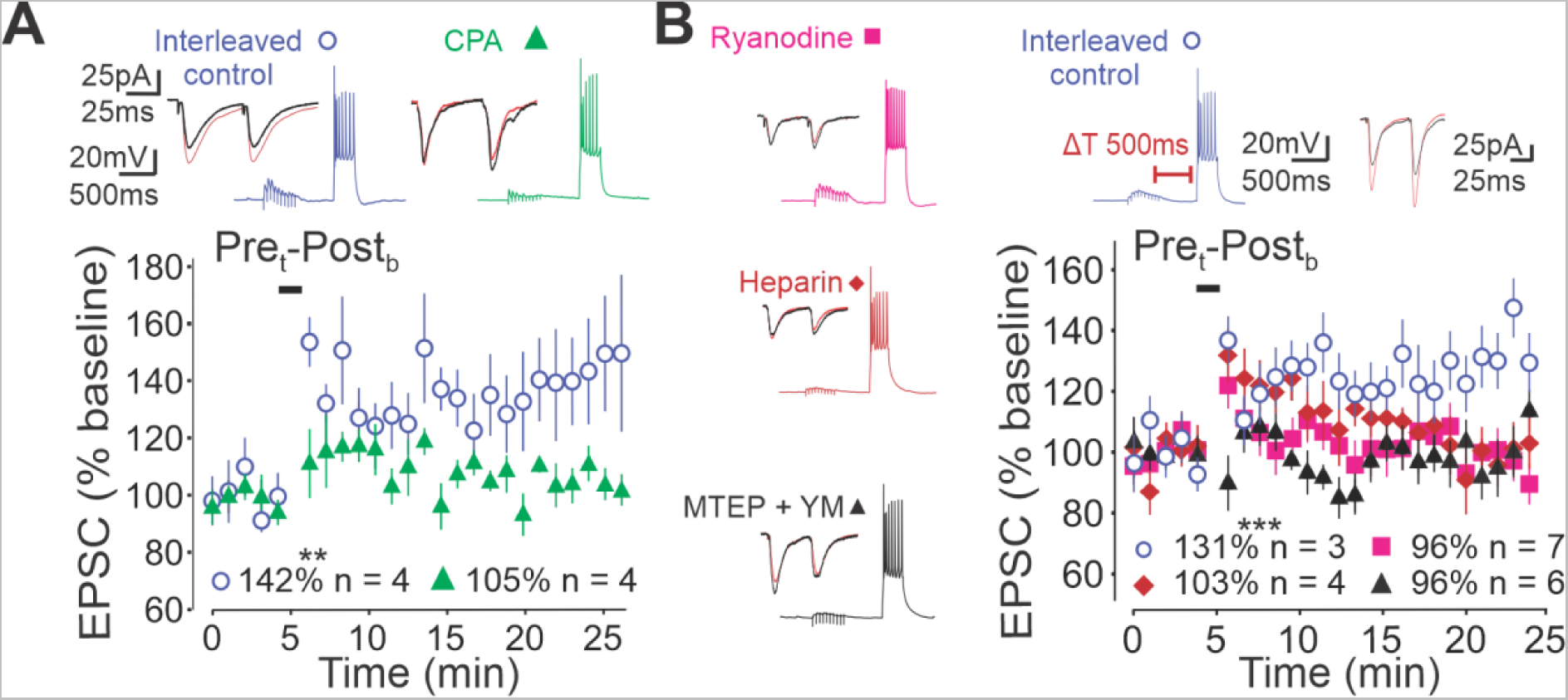
Ca^2+^ stores sustain BTSP in the mPFC. (A) Representative trace of BTSP induction and EPSCs, and resulting normalized EPSC change over time (top) for interleaved control (blue: n = 4 neurons, p = 0.001) and with bath-application of CPA (green: n = 4 neurons, p = 0.19). (B) Representative trace of BTSP induction and EPSCs, and resulting normalized EPSC change over time (top and left) for interleaved control (blue: n = 3 neurons, p = 5.52 x 10^-6^), with intracellular ryanodine (pink: n = 7 neurons, p = 0.47), heparin (red: n = 4 neurons, p = 0.58) or bath-application of MTEP and YM-298198 (black: n = 6 neurons, p = 0.1). **p < 0.01, ***p < 0.001. Data are shown as mean ± sem.

Thus, STAPCD and BTSP in L5 pyramidal neurons abide by similar temporal induction rules and are sustained by an overlapping molecular makeup. Collectively, these results provide convergent evidence for STAPCD as a plausible mechanistic underpinning of BTSP and as a cellular substrate of eligibility traces in cortex. Ergo, STAPCD enacts an eligibility trace that contribute to the binding of temporally separated cues and thus provide an appealing model of associative memory formation.

## Discussion

The high number of paired pre- and postsynaptic stimuli required to induce STDP, along with their strict temporal relationships, renders this plasticity model attractive as a statistical correlation detector during periods of high neural activity, for instance those occurring during early circuit development. This form of plasticity is however less attractive to address the temporal credit assignment problem proper to associative few-shot learning on behavioral timescales. Here, we report in L5 pyramidal neuron of the PFC robust synaptic potentiation following the pairing of only a few pre- and postsynaptic events that are separated by periods of time (0.5 s – 1.0 s) that are substantially longer than neuronal membrane time constants and the rate of glutamate unbinding from NMDARs ^6, 25, 30, 31^. These temporal features alone exclude the involvement of canonical cellular models of Hebbian plasticity and entail the presence of a latent, slowly decaying process that can bind temporally discontiguous events in single neurons. Using a mixture of simultaneous 2P glutamate uncaging and Ca^2+^ imaging experiments, we observed and parameterized a form of associative short-term plasticity of Ca^2+^ dynamics mediated by internal ER stores that arise from a time-dependent interplay between synaptic and bAP-mediated Ca^2+^ entry. We provide temporal and pharmacological evidence that suggests that this ER-mediated STAPCD bears causal relationships with the BTSP observed at these neurons. Thus, this time-decaying short-term cellular memory process satisfies the requirements expected of synaptic eligibility traces and thus provides a cellular model for associative memory over behavioral timescales.

In agreement with previous findings in juvenile and adult pyramidal neurons in hippocampus ^32, 33^, the profile of cytosolic Ca^2+^ transients in L5 neurons following entry through NMDARs during synaptic activation (or through VGCC during trains of bAPs) was not appreciably amplified by CICR machinery, and rather seemed slightly constrained by SERCA activity. Our results however show that the contribution of ER stores to cytosolic Ca^2+^ transients is dynamic and depends on the state of the ER. Thus, temporally and spatially constrained synaptic activation of apical, but not basal, L5 dendrites induces a NMDAR-, mGluR-, and RyR/IP_3_R-dependent amplification of the intracellular Ca^2+^ transients triggered by a subsequent burst of bAPs inasmuch as they occur within ∼1 second. One parsimonious model for STAPCD is that the level *per se* of Ca^2+^ in ER stores is dynamically regulated by Ca^2+^ entry through either synaptic or VGCC sources, rendering them transiently competent to support CICR/IICR release by a subsequent instructive event. The similar time course and induction rules, along with a common set of molecular players point towards store-mediated memory of Ca^2+^ dynamics as enacting eligibility traces that are permissive to the induction of BTSP at L5 pyramidal neurons. Future work will be required to address the universality of this mechanism.

The temporal profile of BTSP induction at L5 pyramidal neurons is broadly reminiscent, but not identical, to that observed at CA1 synapses ^17^. While the induction rules of BTSP are symmetrical in both regions (*i.e.,* synaptic potentiation is triggered by temporally discontiguous pairs of pre- and postsynaptic events irrespective of their relative order), those in L5 pyramidal neurons displayed a singular profile where inputs arriving within a short temporal window (100 ms - 300 ms), were not potentiated, but slightly depressed. The simulation from our experimentally parametrized computational model suggests that it likely is a consequence of the non-linearity of Ca^2+^ extrusion mechanism like that of SERCA pump activity previously reported ^34, 43, 44^,. The temporal window for the induction of BTSP may underly a functional significance in the mPFC. In mice, sensorimotor associations require the frontal cortex, in which sensory inputs from recurrent connections to proximal dendrites could see their synaptic weight updated by BTSP instantiated by delayed contextual cues from thalamic inputs on apical dendrites on a timescale of approximately 400 ms to 1 s ^19, 52–54^. Locally instructive cues may arise from the supervisory action of other brain areas, as assumed in computational models of error-driven learning ^55–57^. Alternatively, locally instructive cues may be features of the late-stage dynamics of response, as assumed in models of self-supervised learning ^58^. Future work will be required to formally address these possibilities, and to gain a mechanistic understanding of the behavioral manifestations of BTSP in the PFC.

Experimental evidence that indirectly support the biological existence of eligibility traces came into light only relatively recently and largely rest on two distinct set of experimental evidence: 1) the finding that subthreshold STDP-like protocols are rendered effective when supplemented with a delayed instructive cue in the form of a neuromodulatory input ^12, 13, 59^ and; 2) BTSP, where the pairing of temporally discontiguous pre- and postsynaptic activation induces synaptic potentiation ^17^. In the former case, the genesis of eligibility traces *per se* is conjunctive in nature since it relies on both pre- and postsynaptic events, a hallmark of STDP protocols. While BTSP and STAPCD are themselves manifestly conjunctive (*i.e.,* none are triggered by pre-or post-synaptic events alone), the underlying eligibility traces are not. Indeed, subthreshold synaptic activation (*i.e.,* that are calibrated such that it does not elicit postsynaptic spiking) can lead to robust synaptic potentiation if followed by delayed instructive postsynaptic bursting (Fig. 1; or plateau potential ^17, 19^). Admittedly, since the eligibility traces triggered in these conditions rely on activation of NMDARs (and mGluRs), they may be considered as requiring the conjunctive action of synaptic release with a postsynaptic event (*eg.,* a locally induced but poorly propagating dendritic spike), and thus loosely qualify as conjunctive. However, eligibility traces can be induced by trains of postsynaptic spiking alone (Fig. 1F; Fig. 3B) and is therefore manifestly not relying on a conjunctive pre- and postsynaptic operation. In this post/pre implementation, a burst of bAPs thus appears to render a large population of apical synapses eligible for subsequent potentiation, wherein it is the delayed instructive synaptic inputs that are providing synapse specificity for potentiation. Collectively, the plasticity rules involving ER-mediated eligibility traces are fundamentally distinct from those of STDP not only in providing means for how the brain addresses the temporal credit assignment problem, but also by formally identifying combinatorial relationships between firing patterns, temporal coincidence and the synaptic updating that gives rise to learning.

## Acknowledgements

This work was supported by grants from the Fonds de Recherche du Québec (L.C.B.), Natural Sciences and Engineering Research Council of Canada (L.C.B.; J.C.B.), Canadian Institute for Health Research (J.C.B.; R.N.), the uOttawa Brain and Mind Research Institute (L.C.B.; J.C.B.), Brain Canada (J.C.B.), and the Canadian Foundation for Innovation (J.C.B.). We thank members of Drs. Jean-Claude Béïque, Richard Naud and Leonard Maler labs for helpful discussions, and a special thank you to Michael B. Lynn and Dr. Mark Harnett for critically reading our manuscript.

## Author contribution

L.C.B. and J.C.B. conceptualized and designed the experimental research. L.C.B. acquired and analyzed the experimental data. L.C.B. and R.N. designed the computational model. L.C.B. implemented the computational model. L.C.B., R.N. and J.C.B. interpreted the data and wrote the manuscript.

## Data/code availability statement

The datasets generated during the current study are available from the corresponding author on reasonable request. / The codes generated during the current study are available from the corresponding author on reasonable request.

## SUPPLEMENTARY MATERIAL

## Methods

### Experiments

#### Slice Preparation

Acute brain slices from the PFC of C57BL/6 mice P15 – P16 (STDP experiments) and P16 - P36 (BTSP experiments) were prepared in accordance with the University of Ottawa Animal Care Committee. Mice were obtained from The Jackson Laboratory and had access to food and water *ad libitum*. Acute slices were prepared following similar guidelines as previously described ^55^. In brief, mice were anesthetized by inhalation of isoflurane (Baxter Corporation, Canada) and sacrificed by decapitation. The brain was immediately removed and slices were transferred to an ice-cold cutting solution containing (in mM): 119 choline-Cl, 2.5 KCl, 1 CaCl_2_, 4.3 MgSO_4_-7H_2_O, 1 NaH_2_PO_4_, 1.3 sodium L-ascorbate, 26.2 NaHCO_3_, and 11 glucose, and equilibrated with 95 % O_2_, 5 % CO_2_ gas. Slices were sectioned and transferred to a recovery chamber containing artificial cerebrospinal fluid (Ringer) solution (in mM): 119 NaCl, 2.5 CaCl_2_, 1.3 MgSO_4_-7H_2_O, 1 NaH_2_PO_4_, 26.2 NaHCO_3_, and 11 glucose, at a temperature of 37 °C, bubbled with 95 % O_2_, 5 % CO_2_. Slices were left in the recovery chamber for at least 1 hour prior to recordings at room temperature.

#### Whole-Cell Electrophysiology

L5 pyramidal neurons of the mPFC were visualized using an Olympus BX51W1 microscope with submersive LUMplanFL N 40x/0.8W objective. Slices were continuously perfused with Ringer solution (described in *Slice Preparation* section of the Methods). Whole-cell recordings were performed using borosilicate glass patch electrodes (4-6 MΩ; Sutter Instruments, Florida) pulled on a Narishige PC-10 pipette puller (Narishige, Japan), filled with a K^+^-Gluconate-based intracellular solution containing (in mM): 115 potassium gluconate, 20 KCl, 10 sodium phosphocreatine, 10 HEPES, 4 ATP(Mg^2+^), and 0.5 GTP (pH adjusted with KOH); or if otherwise specified, with a cesium internal solution containing (in mM): 115 cesium methane- sulfonate, 5 tetraethylammonium-Cl, 10 sodium phosphocreatine, 20 HEPES, 2.8 NaCl, 5 QX- 314, 0.4 EGTA, 3 ATP (Mg^2+^), and 0.5 GTP (pH adjusted with CsOH). Both solutions had pH 7.25 and osmolarity of 280 – 290 mOsmol/L. Internal solutions were prepared in advance and kept at -80 °C until day of experiments. Unless specified otherwise, GABA_A_ receptor mediated currents were blocked with 100µM internal picrotoxin (PTX; Abcam). Whole-cell recordings were carried out using an Axon Multiclamp 700B amplifier, and voltage and current were low- pass filtered at 2 kHz and sampled at 10 or 20 kHz using a Axon Digidata 1440A (or 1550) digitizer. Experiments were performed at room temperature, except where specified otherwise (30 - 32 °C using an in-line bath heater as well as a stage heater). Access resistance was monitored on each sweep using a 200 ms, 5 mV hyperpolarizing pulse, for voltage clamp recordings, or with a 400 ms, -25 pA current injection for current clamp recordings, each induced at least 800 ms prior to electrical synaptic stimulations. Only recordings with steady access resistance (± 30 %) were included. Liquid junction potential was not compensated for. For experiments conducted in current-clamp, small somatic direct current injection was at times used to maintain membrane potential between -65 mV and -75 mV to reduce probability of spiking by synaptic stimulation (see below).

#### Electrical Stimulation of L5 Proximal Synapses and LTP Experiments

Synaptic inputs were stimulated with a stimulating electrode proximal to L5 neurons to determine the ability of proximal synapses to undergo plasticity. Electrical stimuli of proximal synapses were delivered through a stimulation electrode (borosilicate glass; 4-6 MΩ), filled with Ringer, located < ∼200 µm from the soma of the recorded neuron. The stimulating electrode was controlled with an iso-flex ULC stimulation box and stimuli were 0.1 ms in duration. The amplitude of electrical stimulation were experimentally adjusted so that to induce EPSCs of ∼20 pA - 300 pA. For most plasticity experiments, two EPSCs were induced at 50 ms interval in voltage clamp every 20 s for 10 minutes (baseline recordings) and 20 minutes (post-induction recordings). The induction protocol for BTSP experiments were carried out in current clamp mode and consisted of a train of 10 (pre_t_) electrical stimulation at 20Hz (alone or) paired with a 300 ms 400 pA - 600 pA (post_b_) current injection, at variable delays. The BTSP pairing protocol was repeated 5 times every 15 seconds. STDP pairings consisted in a single presynaptic stimulation paired with a 50 ms 300 pA current injection induced 120 times at 33 Hz.

#### Data Analysis - Electrophysiology

Electrophysiological recordings were analyzed on Clampfit 10.7 (Molecular Devices), and Python programming language (Numpy and Scipy libraries). n refers to the number of cells recorded per experiments and all experiments were conducted on different neurons unless stated otherwise. The first EPSC of the paired-pulse in the baseline and post-induction stimulations were used for plasticity analyses. The magnitude of potentiation/depression was calculated as the mean percent change in EPSC amplitude between baseline and the last 5 minutes of post-induction recordings. Cells whose synaptic weight changed more than 3 times the standard deviation away from the mean were removed from analyses. The amplitude of the first and second EPSC in the paired pulse was used for PPR analysis. The PPR was calculated as the mean change in amplitude of the second EPSC compared to the first for baseline and post-induction recordings (post-induction/baseline).

#### Two-Photon Microscopy

For Ca^2+^ imaging experiments, 20 µM Alexa Fluor 594 hydrazide (Thermo Fisher Scientific) and 0.2 mM Fluo-4FF (pentapotassium salt; Thermo Fisher Scientific) were added to the recording electrode to visualize morphology and intracellular Ca^2+^, respectively. For glutamate uncaging experiments, 2.5 mM 4-Methoxy-7-NitroIndolinyl (MNI)-caged-glutamate- trifluoroacetate (Femtonics) was added to the extracellular solution. Two-photon imaging and glutamate uncaging were performed simultaneously using two Ti:Sapphire pulsed lasers (MaiTai DeepSee, Spectra Physics), with the imaging laser set to 810 nm (Alexa Fluor 594 and Fluo-4FF) and the uncaging laser set to 720 nm (for glutamate uncaging). Independent acousto-optic modulators were coupled to a dual galvanometer scanning system (Olympus MPE-1000; BX61WI upright microscope) with a LUMplanFL N 60x/1.0W lens. Image acquisition and stimulation patterns were controlled using Olympus FV10-ASW software (Version 3). Neurons were imaged >10 min after break-in. Proximal apical dendrites were surveyed to find a dendritic segment within the same XY plane containing at least 5-6 spines. Electrophysiological recordings were performed as described in the *Whole-Cell Electrophysiology* section of the Methods. In voltage-clamp at a holding potential of -70 mV, uncaging stimulations were tuned to evoke small EPSCs (2 pA - 20 pA) for each stimulated spine, for a resulting compound EPSC of ∼ 10 pA - 120 pA for clustered activation. The stimulations were subsequently delivered in current clamp for a resulting compound EPSP of ∼ 0.1 mV - 5 mV. These stimulations induced at 20Hz for 500ms, either on their own, or paired with a 300 ms 400 pA - 600 pA somatic current injection. Ca^2+^ imaging was performed simultaneously with uncaging with a line scan probing individual dendritic spines and parent dendrite at ∼0.66 kHz (∼1.5ms/line). Kalman averaged (2 frames) images were obtained to visualize spine morphology. Z-stacks along the entire apical dendrites were obtained to visualize whole-cell morphology.

#### Data Analysis - Two-Photon Microscopy

Ca^2+^ signals were analyzed through Python programming language. Ca^2+^ signals were filtered using a smoothing filter based on the convolution of the signal with a scaled window (Hanning function). Uncaging artifacts were removed to analyze synaptic Ca^2+^ signals. Experiments where Ca^2+^ fluorescence exhibited anomalous behavior (*i.e.,* high baseline Ca^2+^ fluorescence, failure to decay or goodness of fit (R^2^) < 0.4) were excluded from analyses to ensure that recordings were restricted to healthy dendrites. We also excluded experiments where the Ca^2+^ signals induced by the baseline burst of bAPs (*i.e.,* ‘I’) differed from the recovery burst of bAPs (*i.e.,* ‘III’) by more than 35 %, which we inferred reflected poor dendritic health. To quantify Δ[Ca^2+^], we computed ΔF/F_o_ to minimize signal-noise variability and to facilitate comparison between neurons. For each scan, the mean baseline fluorescence (F_o_) was subtracted from the Ca^2+^ signal (F), which was subsequently divided by the mean baseline fluorescence (F_o_): ΔF/F_o_ = (F – F_o_) / F_o_. F_o_ was calculated using the average fluorescence over the 20 linescans just prior to stimulation. At times, during paired stimulations of presynaptic and postsynaptic spiking, the Ca^2+^ signal failed to completely decay by the time of the second stimulation. Thus, we calculated a mono-exponential fit of Ca^2+^ decay to estimate the baseline (F_o_) at the cross-section between the fit and the peak amplitude of the paired Ca^2+^ signal. In case of poor fitting, the baseline fluorescence (F_o_) was overshooted by measuring the Ca^2+^ fluorescence at the start of the second (paired) stimulation. The decay time constant (τ) was calculated from a mono-exponential fit of Ca^2+^ signal from the peak fluorescence to the end of the stimulation. Synaptic Ca^2+^ events with ΔF/F < 0.1 were discarded from analyses except for synaptic Ca^2+^ during D-AP5 experiments. For distance dependence experiments, a synaptic ΔF/F > 0.1 was measured in the synaptic cluster before recordings outside of the synaptic cluster. Fits were obtained using the curve_fit method from the Scipy library. Images of entire neurons were analyzed on ImageJ software and assembled on Paint.

#### Statistics

Normality of data was determined using a Shapiro wilks test on SciPy Stats package in Python. Un-normalized EPSC data (plasticity experiments) followed a normal distribution p > 0.05. As such two-sided paired or unpaired Student t-test were used in plasticity experiments to compare before and after EPSCs, and between group data, respectively (p < 0.05 indicated with an asterisk (*) to determine statistical significance). Decay time constant and ΔF/F data did not follow a normal distribution (p < 0.05), therefore, we used two-sided Mann-Whitney U to compare groups and the two-sided Wilcoxon Signed Rank Test to compare before and after experiments. We used a Bonferroni correction to correct for multiple comparisons where appropriate. Pearson correlation tests were used in correlational analysis. Data are presented as means ± SEM.

#### Drugs and Chemicals

Drugs were added as described in text. Concentrations used for drugs not described elsewhere for the internal solution were: 20 mM 1,2-bis(o-aminophenoxy)ethane-N,N,N′,N′-tetraacetic acid (BAPTA; tetrapotassium salt); 1 mg/ml Heparin (Sodium salt); 100 µM Ryanodine, and for external solution were: 30 µM Cyclopiazonic Acid (CPA); 100 µM 5-phosphono-D-norvaline (D-AP5); 10 µM 3-[(2-methyl-1,3-thiazol-4-yl)ethynyl]-pyridine (MTEP); 5 µM YM298198 (hydrochloride), 20 µM 6-cyano-7-nitroquinoxaline-2,3-dione (CNQX). All reagents were bought from Abcam (Cambridge, MA).

### Model

#### Model - Ca^2+^ Dynamics

Ca^2+^ dynamics emerging from pre- and postsynaptic spike trains, are defined here as *S_pre_(t)* and *S_pos_(t)*, respectively. We reproduced the presynaptic spiking in our experiments with a spike train consisting of 10 presynaptic spikes at 20 Hz (for pre_t_). S_pos_ (for post_b_) was modeled based on the average number of spikes elicited by the current injection (∼ 8 spikes) during experimental recordings, produced regularly during the 300 ms time frame (28.5 Hz) (see Fig. S2).

Pre- and postsynaptic spikes trigger changes in postsynaptic Ca^2+^ approximated respectively by *I_syn_ = k_syn_ S_pre_* and *I_AP_ = k_AP_ S_pos_*, together giving *I_Ca_ = I_syn_* + *I_AP_*. The temporal dynamics of postsynaptic Ca^2+^ (*c,* baseline corrected) are given by:

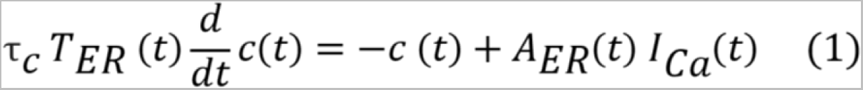

Where *τ_c_* is the decay time constant of cytosolic Ca^2+^ in the absence of ER contributions. *A_ER_* and *T_ER_* model the effect of ER on the amplitude and decay time of Ca^2+^ transients, respectively (see below).

ER Ca^2+^ concentration, *e*, is modeled as being filled at the time of the last event, whether presynaptic (*t_pre_*) or postsynaptic (*t_pos_*). ER Ca^2+^ is assumed to decay exponentially on the time scale of *τ_ER_*:

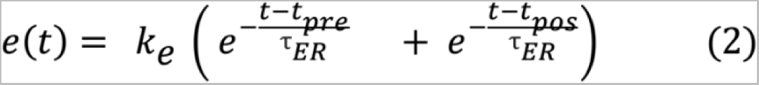

Where *k_e_* is the trace strength upon induction.

The factor *A_ER_* in Eq. (1) modulates the amplitude of cytosolic Ca^2+^ transients. It is made dynamic so as to give larger amplitudes when the ER is filled. It is modeled as a readout of the state of ER at the time of an instructive signal *t_*_* with *f(t_*_) = k_f_ e^-t^_*_^/τf^* representing an exponentially decaying function with an amplitude of *k_f_*, and decay time constant *τ_f_*, that accelerates the effect of decaying *e* in time.

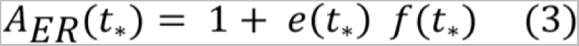

Similarly, the factor *T_ER_* modulates the decay time of postsynaptic Ca^2+^ based on a readout of ER state, and a dynamic state *r*.

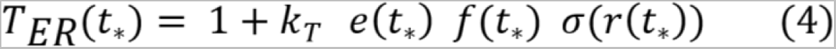

where σ is the sigmoid function of *r* with sensitivity *β*, and *k_t_* is a strength modulating factor. The dynamic state *r* relates to the Ca^2+^ dependence of SERCA pumps rate of transport ^43, 44^. It was introduced to match the observed refractory period. The relative refractory period is modeled as being triggered by previous increases in cytosolic Ca^2+^:

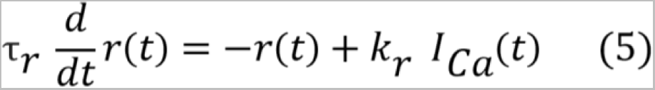

Where *k_r_* controls the coupling between cytosolic Ca^2+^ and the refractory state and *τ_r_* is the decay time constant of *r*.

### Model – Fitting

Parameters were fitted using a set of initial parameters and an optimize minimize function using SciPy module in python to obtain the least squared error (*error_c_*) between experimental data and the model parameters using the equations below:

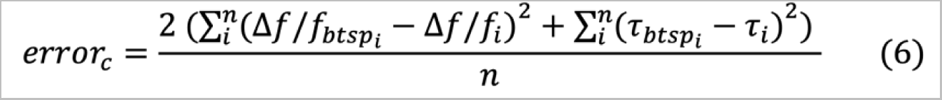

Where *i* is an index labeling each time point tested in the experimental protocol (250 ms - 4.5 s pre_t_-post_b_, 750 ms post_b_-pre_t_), n is the total number of time points across all protocols (n = 7), Δ*F/F_btsp_* and *τ_btsp_* are the experimental normalized peak fluorescence and decay time constant, and ΔF/F and *τ* are the values for the normalized peak Ca^2+^ and decay time constant obtained during parameter fitting for the model.

The decay time constant was obtained using the same fitting method as for the experimental data, except that in experimental recordings the fit was obtained from the peak of Ca^2+^ due to low temporal resolution of the dye, whereas in computational experiments, the decay was fitted from the last spike in the event.

We found a single set of parameters that reproduced our Ca^2+^ data in all three protocols and across all tested timescales (Fig. 5C-K; Table S1).

## Supplementary Figures & Tables

**Table S1.** Fitted Parameters for Ca^2+^ Model.

| Parameters | Definitions | Values |
| --- | --- | --- |
| $k_{\text{syn}}$ | Amplitude of synaptic currents | 5.11 |
| $k_{\text{AP}}$ | Amplitude of AP currents | 4.83 |
| $\tau_c$ (ms) | Decay time constant of $\text{Ca}^{2+}$ | 23.48 |
| $k_e$ | Amplitude of $\text{Ca}^{2+}$ currents in ER | 1.91 |
| $\tau_{\text{ER}}$ (ms) | Decay time constant of ER state | 999.25 |
| $k_f$ | Amplitude of function $f$ | 0.29 |
| $\tau_f$ (ms) | Decay time constant of function $f$ | 1618.17 |
| $k_t$ | Strength modulating factor | 0.91 |
| $\beta$ | Sensitivity of sigmoidal function | 0.0024 |
| $k_r$ | Coupling between cytosolic $\text{Ca}^{2+}$ and $r$ | 80.81 |
| $\tau_r$ (ms) | Decay time constant of $r$ | 664.91 |

**Figure S1.**
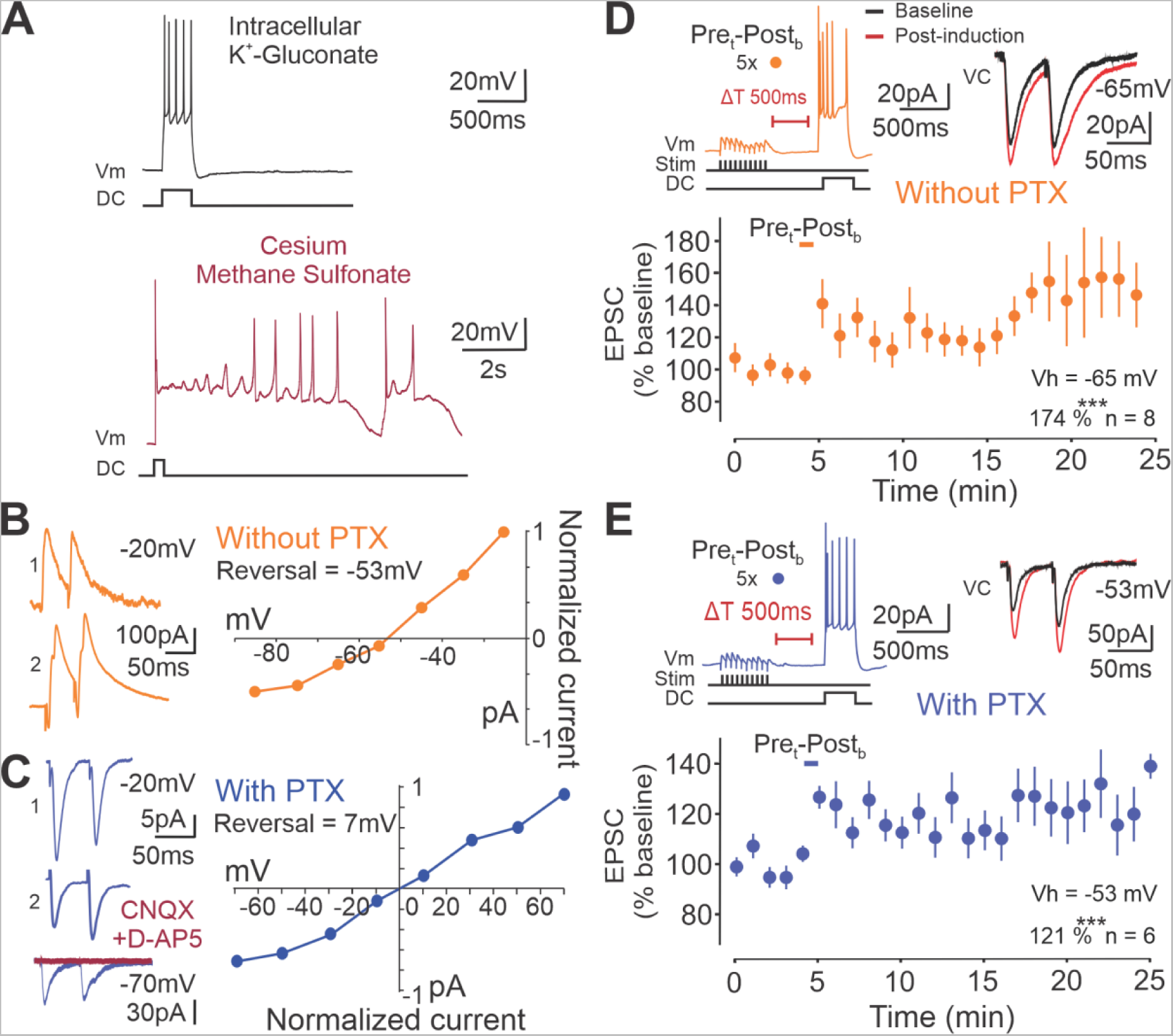
PTX does not affect BTSP. (A) Representative traces of membrane voltage of L5 pyramidal neurons of the mPFC during a post_b_ with K^+^-Gluconate (top) or Cesium (bottom) based intracellular solution. Due to the time for repolarization, we used K^+^-Gluconate internal solution for the remaining of our recordings. (B) Two representative EPSC traces obtained by paired stimulation at 50 ms interval without PTX while holding at -20 mV using K^+^-Gluconate intracellular solution (left). Amplitude of EPSCs (pA) against membrane voltage (mV). (C) Same as B except in the presence of intracellular PTX (blue), and example trace of EPSCs after 10 µM CNQX and 100 µM D-AP5 wash-in (red; lower left). (D) Representative trace of pre_t_- post_b_ pairings and EPSCs recorded without PTX at -65 mV (top) and resulting normalized EPSCs change over time (n = 8 neurons, p = 5.63 x 10^-8^, bottom). (E) Same as in D except with PTX and holding at -53 mV (at reversal; n = 6 neurons, p = 0.00011). Vh: Voltage holding. Data are shown as mean ± sem.

**Figure S2.**
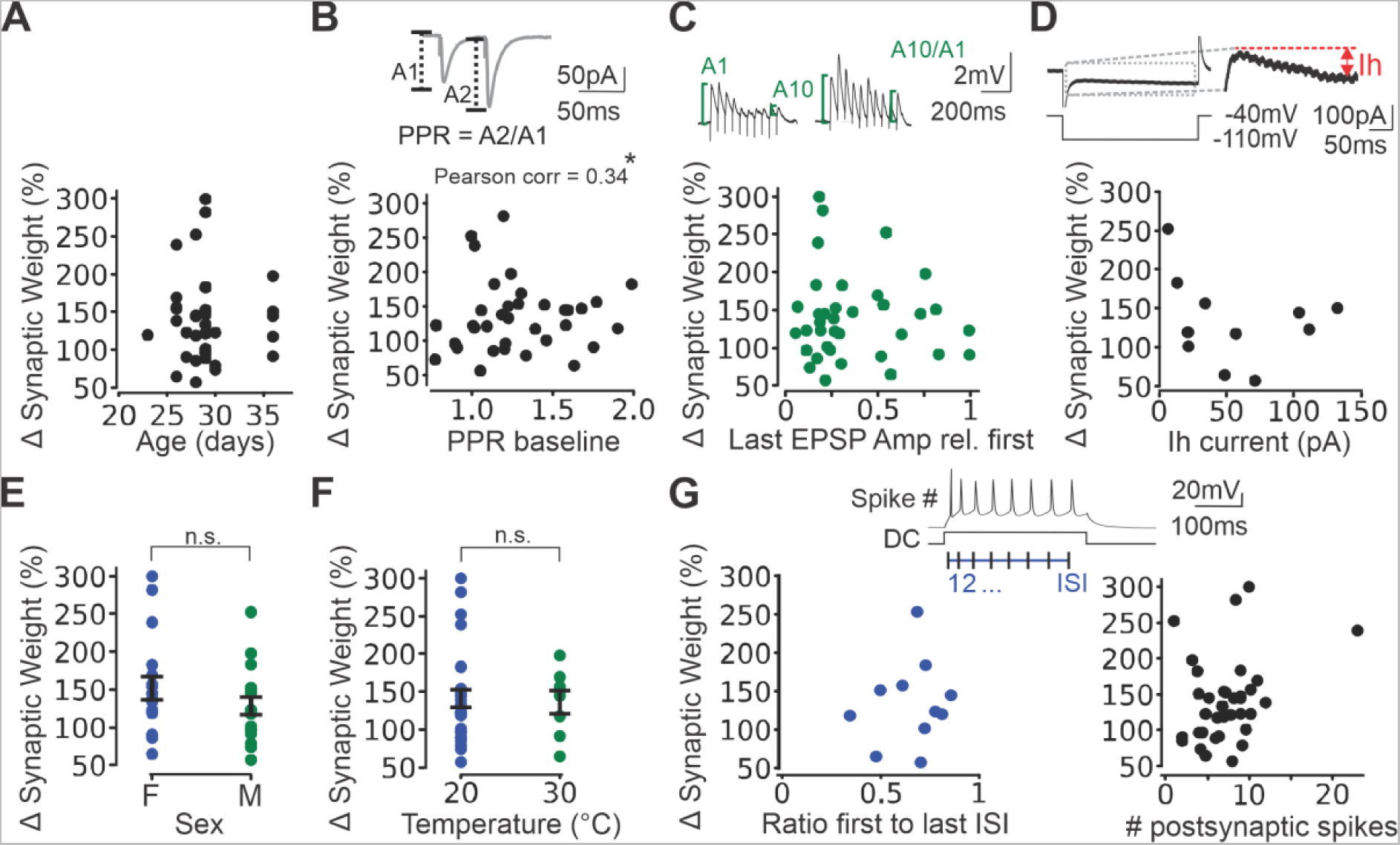
Generalization of synaptic rules during pre_t_-post_b_ and post_b_-pre_t_ pairings (500 ms – 750 ms). (A) LTP magnitude against age in days at time of death (n = 32 neurons, p = 0.94). (B) Representative EPSC trace obtained during baseline recordings (top). LTP magnitude against PPR at baseline (n = 35 neurons, p = 0.04, coefficient = 0.34). (C) Representative trace of EPSPs during pre_t_-post_b_ pairings (top). LTP magnitude against synaptic transmission in a train calculated as the ratio of the last EPSP amplitude (A10) to the first (A1) in the train (n = 36 neurons, p = 0.63, bottom). (D) Representative trace of voltage clamp recordings of a cell depolarized from -40 mV to -110 mV, and schematic of Ih current amplitude measurement (top). LTP magnitude against Ih current (n = 11 neurons, p = 0.43, bottom). (E) LTP magnitude against sex of the animal (female (F): n = 15 neurons, male (M): n = 13 neurons, p = 0.22). (F) LTP magnitude against recording temperature (20: n = 18 neurons, 30: n = 7 neurons, p = 0.84). (G) Representative trace of a post_b_ induced in current clamp and calculation of # of spikes and inter-spike interval (ISI; top). LTP magnitude against the duration of the last ISI to the first (n = 14 neurons, p = 0.69, bottom left). LTP magnitude against # of postsynaptic spikes during post_b_ induced by 400 pA to 600 pA direct current injection (n = 35 neurons, p = 0.08, bottom right). Rel: relative. DC: Direct current. Data are shown as mean ± sem.

**Figure S3.**
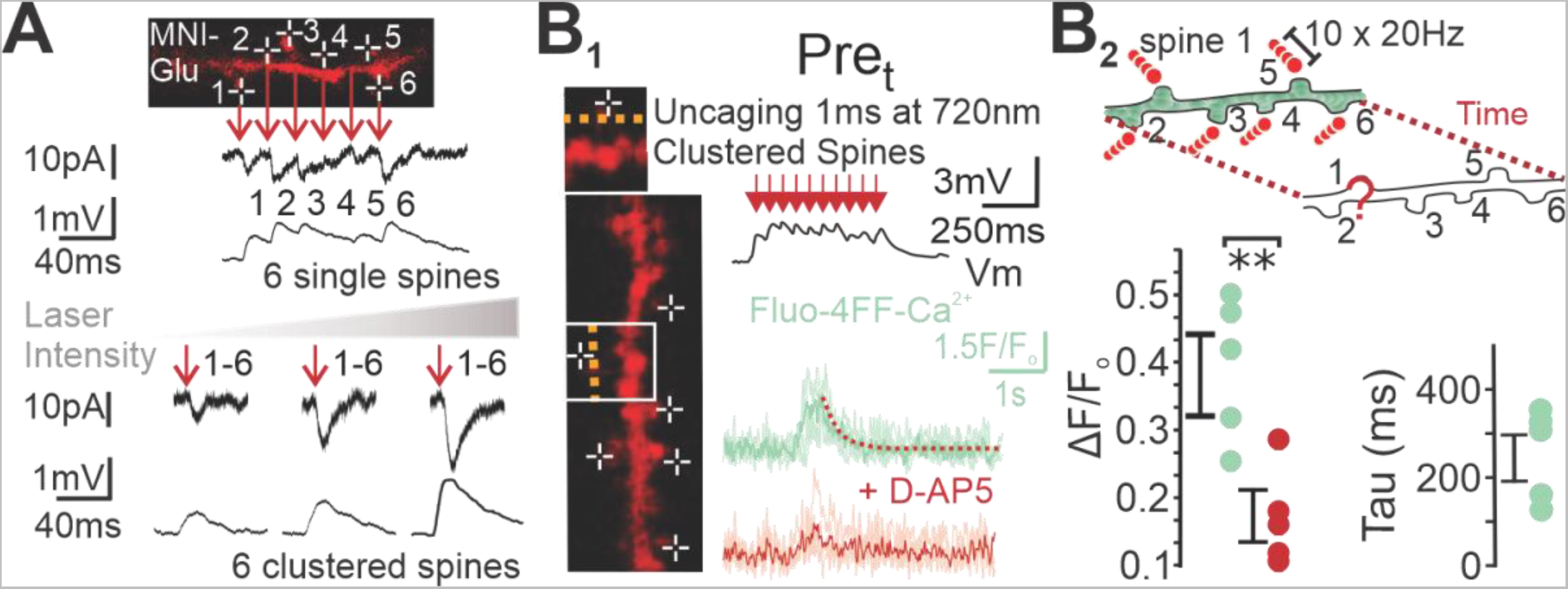
Ca^2+^ dynamics of uPre_t_ induced on clustered spines on proximal dendrites of L5 mPFC pyramidal neurons. A) Example image of proximal apical dendrite of L5 pyramidal neurons on which glutamate uncaging was performed on 6 individual spines at 40% laser power (top), and on spatially and temporally clustered spines (6 spines at the same time) at varying laser powers (20 % - 40 %), with their representative voltage and current clamp traces (bottom). (B) B_1_ Representative trace of uPre_t_ induced with glutamate uncaging on 6 clustered spines of the proximal apical dendrites of L5 pyramidal neurons (top), and resulting Ca^2+^ traces in control condition (green) and with bath-application of D-AP5 (red) (bottom). B_2_ Cartoon of experiment (top) and resulting ΔF/F (p = 0.0012) and tau values (bottom). **p < 0.01; Data are shown as mean ± sem.

**Figure S4.**
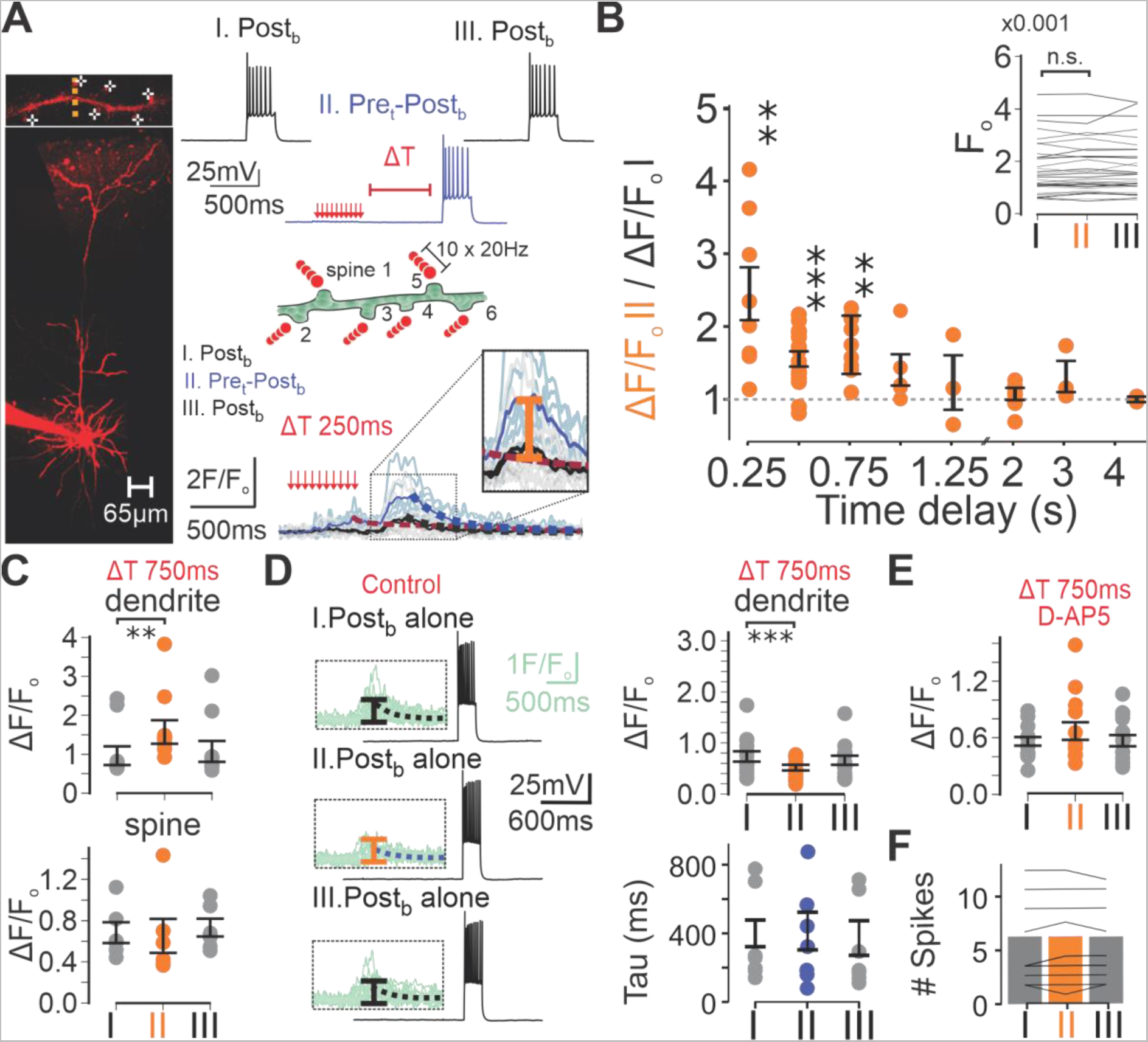
Ca^2+^ dynamics elicited by pre_t_-post_b_ pairings. (A) Example image of L5 pyramidal neuron and proximal apical dendrite on which clustered glutamate uncaging and line scans of the dendritic segment were performed (left). Representative trace of Ca^2+^ during three post_b_ induced at 30 s interval with the second post_b_ preceded by pre_t_ at varying time interval (top right). Cartoon of synaptic stimulation (middle right). Ca^2+^ fluorescence recordings with Fluo- 4FF in proximal apical dendrites of L5 pyramidal neurons for 250 ms delay protocol (bottom right). (B) Normalized relative ΔF/F values of post_b_ preceded by pre_t_ compared to the first post_b_ given alone as shown in A (250 ms: n = 8 dendrites, p = 0.0078, 500 ms: n = 15 dendrites, p = 0.00026, 750 ms: n = 9 dendrites, p = 0.002, 1000 ms: n = 5 dendrites, p = 0.13, 1250 ms: n = 3 dendrites, p = 0.75, 2000 ms: n = 8 dendrites, p = 0.546, 3000 ms: n = 3 dendrites, p = 0.25, 4500 ms: n = 3 dendrites, p = 1.0). Baseline fluorescence (F_o_) prior to each of the three post_b_ shown in A across all time delays (inset, n = 47 dendrites, p = 0.30). (C) ΔF/F values for protocol shown in A for time delay of 750 ms in dendrite (same as in B) and spine (n = 6 spines, p = 0.84). (D) Representative trace of three post_b_ given at 30 s interval without pre_t_ and resulting Ca^2+^ trace (left). Dendritic ΔF/F (n = 14 dendrites, p = 0.00061) and tau (n = 7 dendrites, p = 0.38) values (right). (E) ΔF/F values for post_b_ Ca^2+^ events from protocol shown in A at 750 ms delay with the presence of D-AP5 (n = 12 dendrites, p = 0.27). (F) Number (#) of spikes for each post_b_ for dendritic data shown in C (n = 9 dendrites, p = 0.23). **p < 0.01, *** p < 0.001. Data shown as mean ± sem.

**Figure S5.**
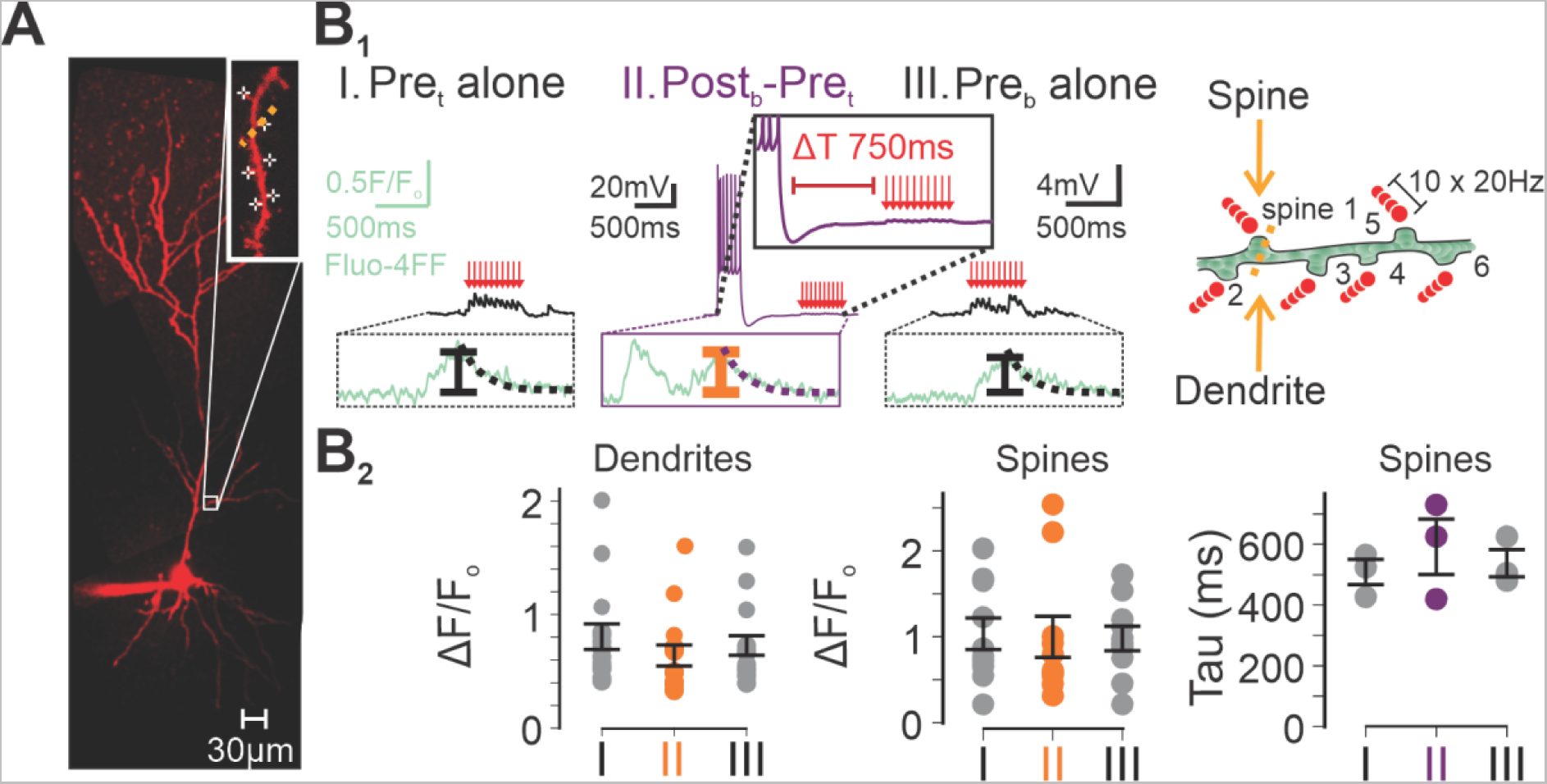
STAPCD is bidirectional. (A) Example image of L5 pyramidal neuron and proximal apical dendrite on which clustered glutamate uncaging and line scans of the dendritic segment were performed. (B) B_1_ Representative voltage and mean Ca^2+^ traces of three pre_t_ induced at 30 s intervals with the second pre_t_ preceded by post_b_ 750 ms prior (left). Cartoon of synaptic stimulation (right). B_2_ Resulting ΔF/F values in dendrites (n = 15 dendrites, p = 0.23, left) and spines (n = 10 spines, p = 0.70, middle) and spine tau values (n = 3 spines, p = 0.5, right). Data shown as mean ± sem.

**Figure S6.**
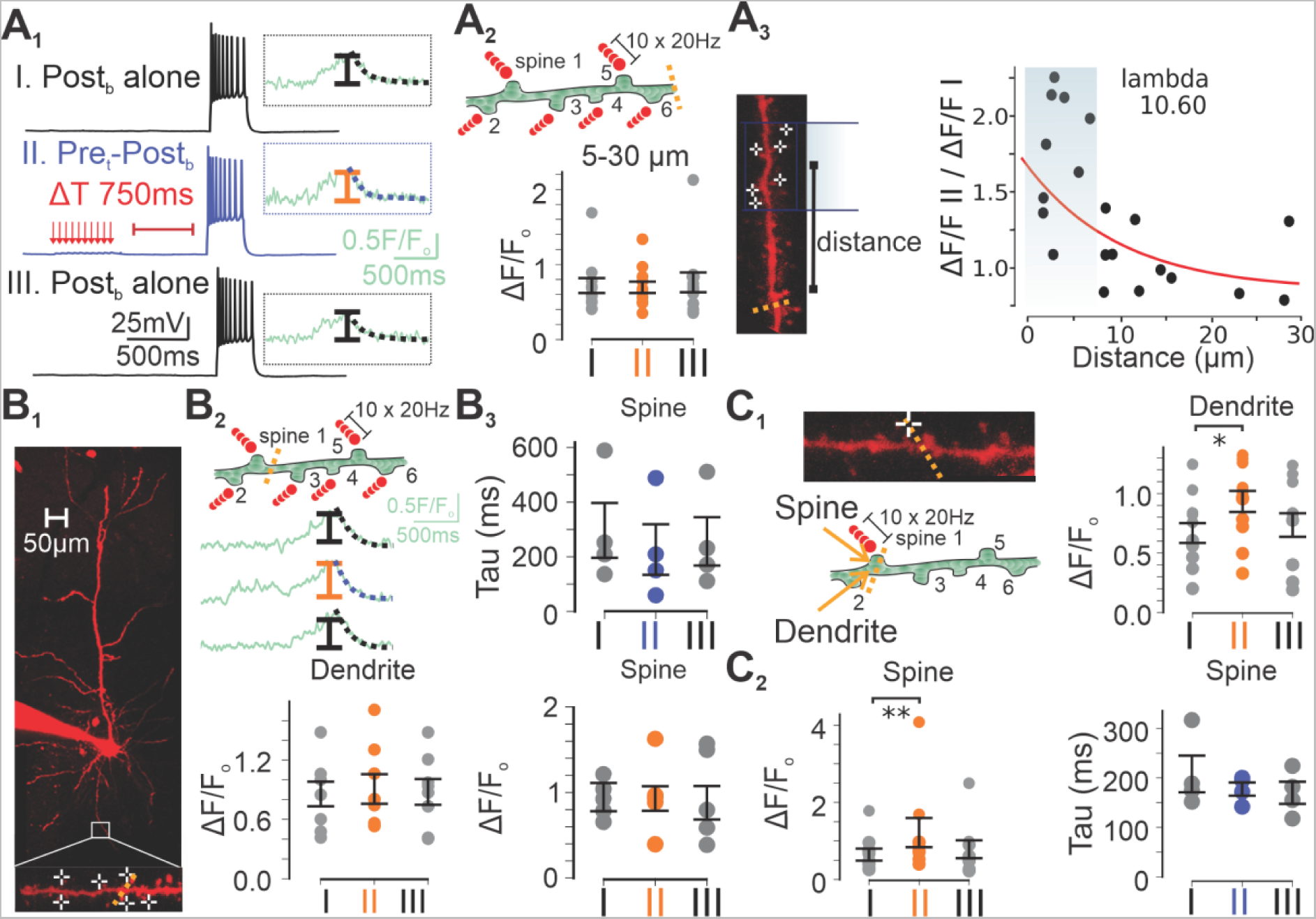
Properties of STAPCD during effective BTSP. (A) A_1_ Representative voltage and mean Ca^2+^ traces of three post_b_ induced at 30 s interval with the second post_b_ preceded by pre_t_ 750 ms prior. A_2_ Cartoon of synaptic stimulation showing line scans performed 5-30 µm away from the middle of the synaptic cluster (top) and resulting dendritic ΔF/F values (n = 11 dendrites, p = 0.64, bottom). A_3_ Example image of L5 pyramidal neuron’s proximal apical dendrite on which clustered glutamate uncaging and line scans of the dendritic segment 5-30 µm away were performed (left). Distance dependence of relative normalized ΔF/F values (n = 20 dendrites, right). (B) B_1_ Example image of L5 pyramidal neuron and proximal basal dendrite on which clustered glutamate uncaging and line scans were performed. B_2_ Cartoon of synaptic stimulation (top), mean Ca^2+^ traces (middle) and resulting dendritic ΔF/F values (n = 7 dendrites, p = 0.81, bottom). B_3_ Resulting spine values of tau (n = 4 spines, p = 0.13, top) and ΔF/F (n = 5 dendrites, p = 0.81, bottom). (C) C_1_ Synaptic stimulation example (left) during protocol shown in A_1_ but on a single spine, and resulting dendritic ΔF/F values (n = 11 dendrites, p = 0.032, right). C_2_ Same as C_1_ (right), except for spines ΔF/F (n = 9 spines, p = 0.0039, left) and spine tau values (n = 4 spines, p = 0.63, right). *p < 0.05, **p < 0.01. Data shown as mean ± sem.

**Figure S7.**
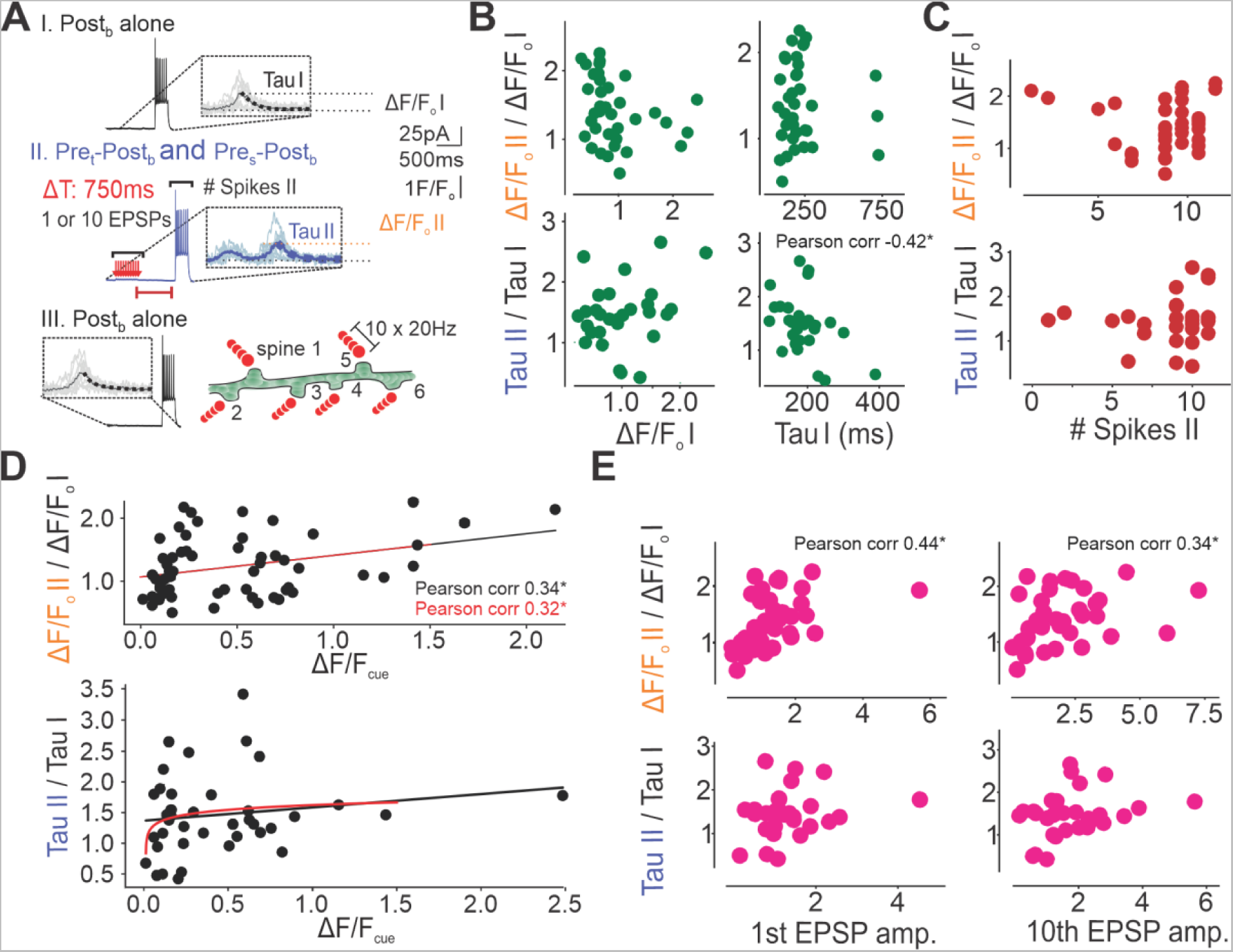
Relationship between STAPCD, Ca^2+^ and voltage. (A) Representative STAPCD protocol. (B) Relative ΔF/F and tau values against ΔF/F and tau value of the first post_b_ (Relative ΔF/F against ΔF/F_1_: n = 36 dendrites, p = 0.09; Relative ΔF/F against Tau_1_: 36 dendrites, p = 1.0; Relative Tau against ΔF/F_1_: n = 29 dendrites, p = 0.16; Relative Tau against Tau_1_: n = 29 dendrites, p = 0.02, correlation coefficient = -0.42). (C) Relative ΔF/F and tau values against # spikes in the second post_b_ (ΔF/F: n = 36 dendrites, p = 0.69; Tau: n =29 dendrites, p = 0.48). (D) Correlation between relative ΔF/F (top; n = 57 dendrites, linear black (p = 0.01, correlation coefficient = 0.34) and logarithmic red (p = 0.02, correlation coefficient = 0.32)) and tau (bottom; n = 40 dendrites, linear (black, p = 0.32) and logarithmic (red, p = 0.19)) and ΔF/F from the cue preceding the paired input (pre_t_ during pre_t_-post_b_, or post_b_ during post_b_-pre_t_,). (E) Relative ΔF/F and tau values against the 1^st^ and last EPSP in the pre_t_ (Relative ΔF/F against 1^st^ EPSP: n = 36 dendrites, p = 0.01, correlation coefficient = 0.44; Relative ΔF/F against 10^th^ EPSP: n = 36 dendrites, p = 0.04, correlation coefficient = 0.34; Relative tau against 1^st^ EPSP: n = 29 dendrites, p = 0.25; Relative tau against 10^th^ EPSP: n = 29 dendrites, p = 0.12). *p < 0.05. Data for B, C and E from protocols pre_t_-post_b_ 500 ms and 750 ms, Data for D same as B, C and E, with the addition of post_b_-pre_t_ 750 ms.

**Figure S8.**
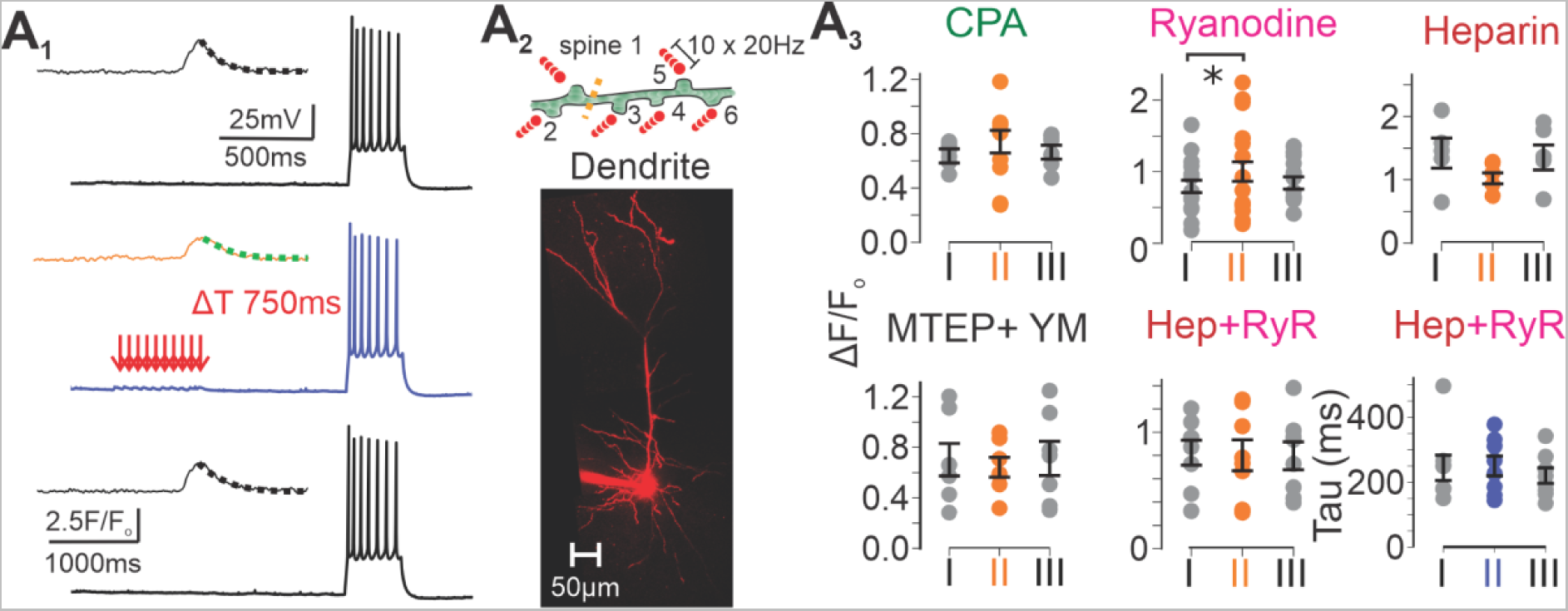
Dependence of STAPCD to ER. (A) A_1_ Representative voltage and mean Ca^2+^ traces of three post_b_ induced at 30 s interval with the second post_b_ preceded by pre_t_ 750 ms prior during bath-application of drugs. A_2_ Cartoon of synaptic stimulation and dendritic line scans (top) and example image of L5 pyramidal neuron (bottom). A_3_ ΔF/F values of protocol shown in A_1_ for cells exposed to either extracellular CPA (n = 8 dendrites, p = 0.31), MTEP and YM298198 (n = 7 dendrites, p = 0.94), or intracellular ryanodine (n = 13 dendrites, p = 0.040) or heparin (n = 5 dendrites, p = 0.13) or a mix of ryanodine and heparin (ΔF/F: n = 5 dendrites, p = 0.44 ; τ: n = 6 dendrites, p = 0.84). Data shown as mean ± sem.

